# Single-cell gene expression prediction from DNA sequence at large contexts

**DOI:** 10.1101/2023.07.26.550634

**Authors:** Ron Schwessinger, Jacob Deasy, Rob T. Woodruff, Stephen Young, Kim M. Branson

## Abstract

Human genetic variants impacting traits such as disease susceptibility frequently act through modulation of gene expression in a highly cell-type-specific manner. Computational models capable of predicting gene expression directly from DNA sequence can assist in the interpretation of expression-modulating variants, and machine learning models now operate at the large sequence contexts required for capturing long-range human transcriptional regulation. However, existing predictors have focused on bulk transcriptional measurements where gene expression heterogeneity can be drowned out in broadly defined cell types. Here, we use a transfer learning framework, seq2cells, leveraging a pre-trained epigenome model for gene expression prediction from large sequence contexts at single-cell resolution. We show that seq2cells captures cell-specific gene expression beyond the resolution of pseudo-bulked data. Using seq2cells for variant effect prediction reveals heterogeneity within annotated cell types and enables *in silico* transfer of variant effects between cell populations. We demonstrate the challenges and value of gene expression and variant effect prediction at single-cell resolution, and offer a path to the interpretation of genomic variation at uncompromising resolution and scale.

## Introduction

Predicting gene expression from DNA sequence remains a challenging task for machine learning[1–5]. Models for mammalian transcriptional regulation need to capture complex sequence logic and interactions over large genomic distances. Solving the expression problem offers a direct path to predicting the impact of genetic variants on transcript levels, a proxy for cellular function. However, established sequence-to-expression models[2–5] use bulk data of broadly defined tissues and cell types. This limits their ability to characterize heterogeneous tissues and pinpoint cell-type-specific variant effects.

Single-cell RNA-seq (scRNA-seq) technology offers a powerful approach to tackle heterogenous tissues beyond the resolution of annotated cell types (Figure 1) but comes at the price of data sparsity. Single-cell RNA-seq experiments frequently lead to more than 90 % zero entries in the cell-by-gene expression matrix[6]. Moreover, the number of recognized human protein coding genes is about 20 k. This bounds the training dataset size along the genes axis and limits the extent to which the number of training examples can be scaled with model capacity. Sequence-to-expression approaches trying to leverage single-cell data have compromised on resolution and granularity by operating on pseudo-bulked data[7] or binarizing expression[8] and operate with limited sequence context.

**Figure 1:**
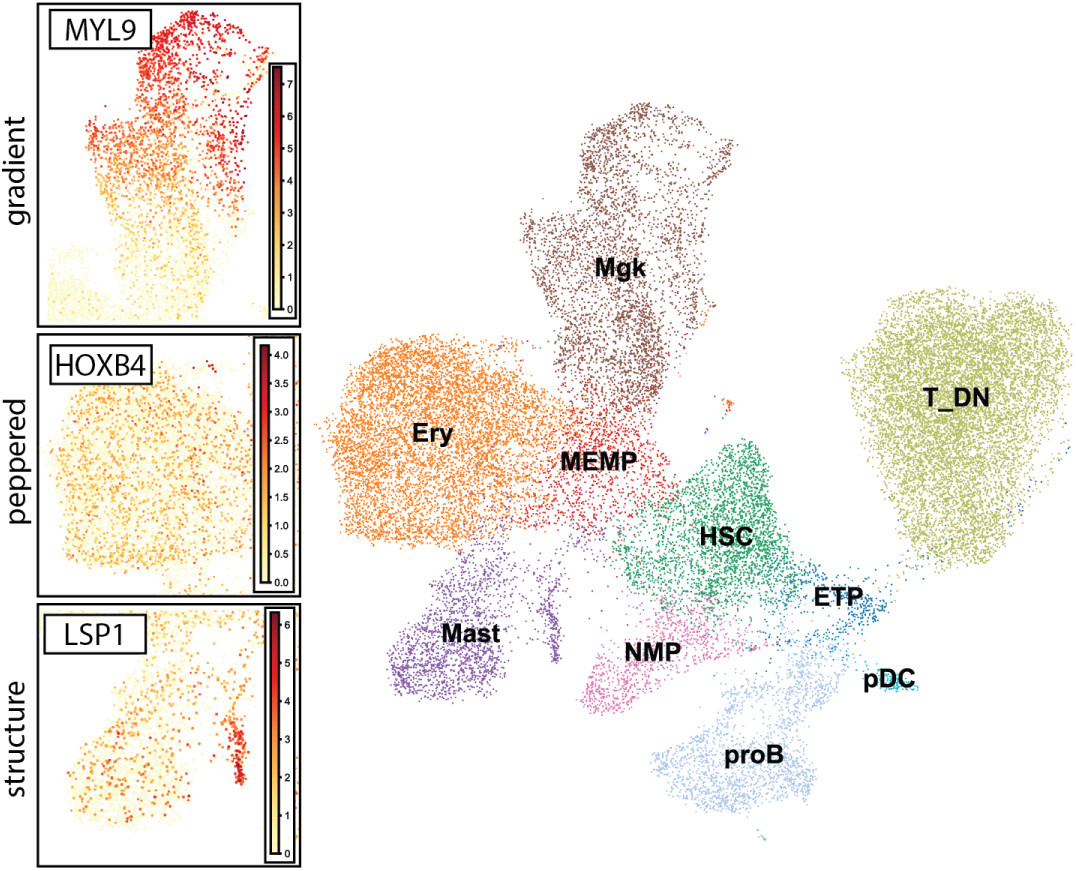
UMAP of a hematopoietic differentiation single-cell dataset [12]. Shown are examples of single-cell gene expression with different, within-cell-type structure that may lead to similar pseudo-bulk expression values: gradient of expression within a cell type (MYL9); peppered but even expression across a cell type (HOXB4); structure within an annotated cell type (LSP1). Cells are colored based on the author provided cell annotation. The color gradient indicates the observed expression per cell. Abbreviations: Ery – erythrocyte; ETP - early thymic progenitor; HSC - Hematopoietic stem cell; Mast - Mast cells; MEMP – megakaryocyte erythrocyte mast cell progenitor; Mgk – megakaryocyte; NMP - neutrophil-myeloid progenitor; pDC - plasmacytoid dendritic cells; proB – pro-B cell. T_DN - double negative thymocyte;

Models pretrained on generic auxiliary tasks followed by transfer learning offer a way to overcome the limitations posed by the finite number of genes. Yet, most self-supervised DNA language models are currently unable to operate at the sequence context necessary to capture human gene regulation[9–11]. In contrast, Enformer[5] uses convolutions for efficient context aggregation. This sacrifices base pair resolution but allows sequence contexts of close to 200 kb. Enformer is trained on a supervised epigenomic prediction task, to predict the building blocks and markers of molecular processes that govern gene regulation. This approach is closer to learning “meaning” in the sense of a natural language task than masked DNA base pair predictions.

In this work, we utilize Enformer as a pre-trained epigenomic model, to create gene embeddings that capture the sequence logic of transcriptional regulation. We demonstrate how to use transfer learning to predict gene expression at single-cell resolution from large scale sequence context. We show that single-cell gene expression models generalize over unseen genes and capture cell-specific gene expression beyond pseudo-bulk resolution. Single-cell gene expression prediction enables variant effect prediction at single-cell and single-base-pair resolution. This allows us to pinpoint variant effects to cells rather than cell types. Characterizing the heterogeneity of variant effect aids us in cell type annotation and in interpreting genetic variation. Scaling the approach up to 650 k cells allows us to predict how variants affecting gene expression in the context of T cell activation act in the context of T cell development.

## Results

### Predicting single-cell gene expression from epigenome model embeddings

We sought to design a flexible, transfer learning framework to predict single-cell gene expression from limited training examples while incorporating large sequence contexts (Figure 2). To this end, we use a DNA sequence model pre-trained on a task with readily-available training examples as a sequence-to-embedding (seq2emb) module and train dedicated, single-cell gene expression modules to map from embeddings to cells (emb2cell). We reasoned that epigenome models serve as good seq2emb modules for gene expression prediction. Recent epigenome models, such as Enformer[5], are trained on a diverse corpus of epigenomic features that include cell type specific CAGE-seq, a technique that maps RNA transcripts roughly to their initiation site and serves as a good proxy for overall RNA transcription levels as measured by RNA-seq[13]. Subsequent work showed an encouraging degree of correlation between Enformer-predicted CAGE-seq tracks with RNA-seq in comparable tissues[14–16]. Enformer is structured into a trunk and head module. The trunk uses convolutional and transformer blocks to embed DNA sequence windows of 128 bp. The head is trained to predict epigenomic features associated with each sequence window. Enformer uses a large sequence context of nearly 200 kb, but all information sharing across sequence windows is achieved in the trunk module. We reasoned that all information required to predict gene expression is therefore encoded in the embeddings of the sequence windows at which CAGE-seq profiles are enriched, predominantly the transcription start site (TSS). Embeddings of the TSS can be viewed as gene embeddings from the perspective of transcriptional regulation.

**Figure 2:**
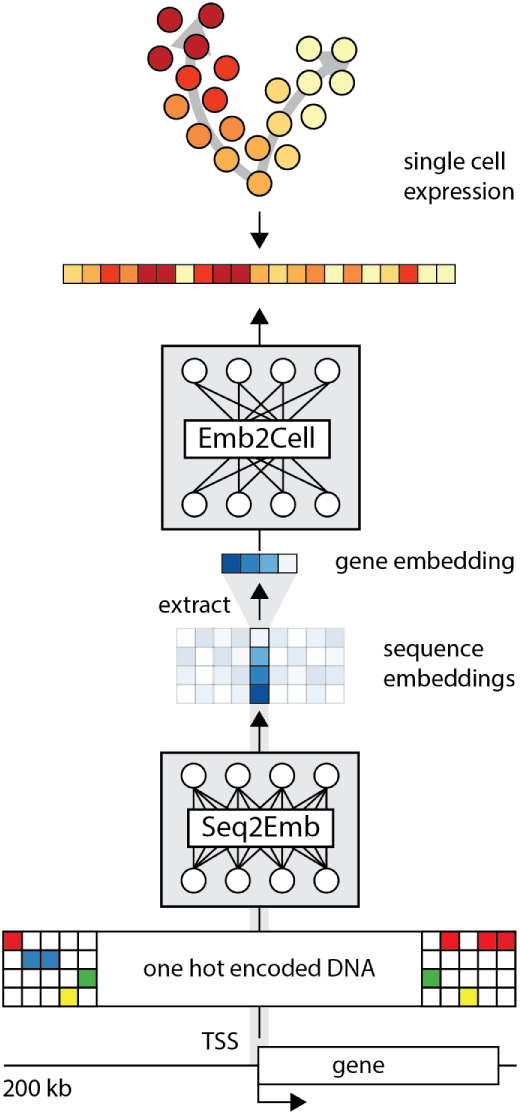
Principal seq2cells design. Every instance is a DNA sequence of ~200 kb centered on a TSS. The one-hot-encoded DNA sequence is processed by a seq2emb module to produce DNA sequence embeddings. The embedding of the TSS is extracted and fed through an emb2cell module to predict the expression of the gene in all cells of a given single-cell dataset. To speed up training, the TSS embeddings of all canonical TSS of protein coding genes are pre-computed.

We use the trunk of Enformer as the seq2emb module (Figure 2). In principle, any DNA sequence to embedding model can be used. For a given gene, the seq2emb module extracts the gene embedding by running a one hot encoded, 200 kb DNA sequence window centered on the TSS through the Enformer trunk. For training speed, we pre-compute the TSS embeddings for all canonical TSS of all protein coding genes (19,986 using Gencode V41[17]). We train an emb2cell module, to predict the single-cell expression of each gene from its gene embedding. Distinct models were trained for each scRNA-seq dataset of interest. We use a two-layer multi-layer perceptron (MLP) as a default emb2cell module that predicts one output per cell. In this formulation, every instance is a gene and the model is trained to predict the cell specific expression. We use validation (1,607) and test (1,930) genes based on the region split used in Basenji2[3] and Enformer[5]. We use a loss function designed to balance between the across cell and across gene correlation (see Methods). We freeze the seq2emb module and find that training saturated after a modest number of epochs (10 to 30).

Appropriate regularization is key since we want to be able to make variant effect predictions on training genes without needing elaborate cross-validation schemes at inference time. In practice, we found that training from epigenome embeddings that have already been trained against bulk cell-type-specific CAGE-seq gene expression easily overfits. Here we use a bottleneck in the MLP, comparatively high dropout rates (up to 0.5) and weight decay to combat overfitting.

At inference time the model can be run using pre-computed embeddings or DNA sequence input. This allows us to predict the effect of sequence variants on single-cell expression by comparing the predicted expression of a reference and variant sequence. We trained single-cell expression models using T cell development data[12], using the hematopoietic stem cell (HSC) focused subset with *~* 30 k cells and the full development dataset of *~* 250 k cells. In addition, we scaled seq2cells to a CD4 T cell activation dataset[18] comprising *~* 650 k cells. We trained models against the log count data after batch correction, as published by the authors. Henceforth we refer to these counts as “raw” counts (see Methods). We focused our evaluations on the smaller dataset (HSC) and explored the scaling potentials and limitations with the full developmental and CD4 T cell activation data.

### Predicting gene expression in single cells is challenging but possible

Sequence-to-expression models are typically evaluated by the correlation of the predicted and observed expression on hold-out genes. The correlation is usually partitioned (Figure 3 A) into: 1) cross-gene, the correlation across genes, averaged over all cells or cell types and 2) cross-cell, the correlation across cells or cell types, averaged over all hold-out genes. The distinction is important because it highlights different aspects of model performance. While cross-gene demonstrates generalizability to unseen genes and hence DNA sequences, cross-cell demonstrates a model’s ability to capture the differences between cells and make cell-(type)-specific predictions. Previous work[3, 5, 14–16] reports both correlations but focusses on discussing the, generally higher, cross-gene performance. However, to achieve highly cell-(type)-specific predictions, the cross-cell performance is key. Model performance can be evaluated at the single-cell level or at the level of pseudo-bulked cell types (Figure 3 B), losing resolution but making it comparable to previous work[5, 7].

**Figure 3:**
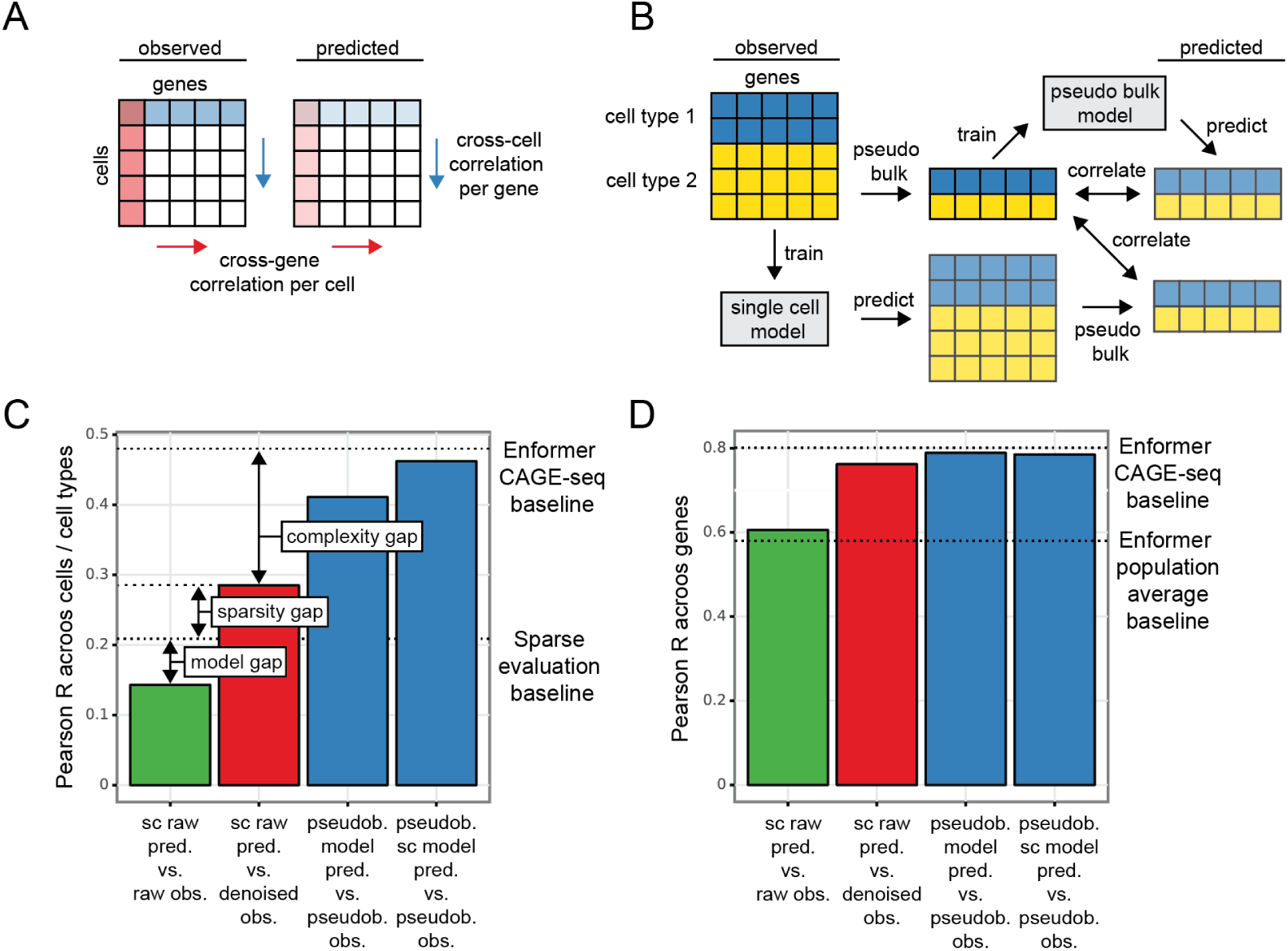
Evaluation scheme and performance of the HSC model. **A)** Observed and predicted single-cell count matrices (cell-by-gene) are compared by calculating the Pearson correlation across cells per gene (blue) and across genes per cell (red). **B)** Pseudo-bulk evaluation scheme: Two modes are compared 1) Pseudo-bulking the observed data into average profiles per annotated cell type; training a model against the pseudo-bulked data; predicting the expression of hold-out genes per cell type and correlating the predicted vs. observed cell-type-specific expression. 2) Training a model against the single-cell data, predicting the expression of hold-out genes in single cells; pseudo-bulking the predictions and correlating the pseudo-bulked predictions with the pseudo-bulked observations. **C)** Mean cross-cell Pearson correlation of single-cell and pseudo-bulk model predictions compared against raw (green), denoised (red) or pseudo-bulked (blue) observations for all test set genes. The Enformer CAGE-seq baseline indicates the mean cross-cell Pearson correlation when finetuning a model on the Enformer CAGE-seq data. The sparse evaluation baseline indicates the mean cross-cell Pearson correlation when assuming denoised observations as ground truth, repeatedly dropping out (set to 0) a number of cells proportional to the gene-specific dropout rate and correlating this sparse observation to the ground truth. Arrows indicate different sources of error: The model gap indicates the distance to a hypothetical perfect model evaluated on the raw observed HSC data . Closing the model gap will increase all single-cell model performance regardless of evaluation. The sparsity gap indicates the drop in performance metric associated with evaluating the smooth predictions against the sparse, raw as opposed to the smooth, denoised counts without changing the actual model performance. The complexity gap indicates the added difficulty of evaluating single-cell expression opposed to pseudo-bulked average expression profiles. **D)** Mean cross-gene Pearson correlation for the HSC dataset. The Enformer CAGE-seq baseline indicates the same as **(C)** but across genes. The Enformer population average baselines indicates the cross-gene Pearson correlation achieved by Sasse et al.[16] when comparing the average cerebral cortex expression of ~ 13 k genes in 839 individuals with the predicted cortex CAGE-seq track of Enformer predicted from the reference genome.

We trained the HSC model on the 30 k cell HSC subset and computed the cross-cell and cross-gene correlation on the hold out test set of 1,930 genes (Figure 3 C & D). We compared the model predictions against the raw observed counts (green bars). Our single-cell model achieved a cross-cell Pearson correlation of 0.143 and cross-gene correlation of 0.606. Because these correlations are substantially lower then results on bulk data[2–5], we next sought to understand the sources of error in and limitations of single-cell gene expression prediction. Single-cell data is sparse. In the HSC dataset, 83 % of entries in the cell-by-gene count matrix are zero. This is a mixture of unexpressed genes and technical dropouts caused by low capturing efficiency in single-cell assays[19]. This causes many observed zeros where read counts are expected. In contrast, the model predicts smooth gene expression values over cells (Figure 4). Correlating a smooth predicted signal with a sparse observed signal yields comparatively low correlation. To estimate expected correlation levels between smooth predictions and sparse observations we need: 1) An understanding of single-cell dropout rates and 2) a ground truth. To understand the influence of single-cell dropout rates we build on the insights from Kim et al.[20]. The rates at which entries in the cell-by-gene matrix are zero are gene-specific and decrease with increasing average expression levels across cells (Supplementary Figure 1 A). Given a ground truth we can mimic the sparse single-cell data by setting to zero entries in the cell-by-gene matrix with probabilities equal to the gene-specific dropout rates. We can repeat this process to estimate a per gene and an overall expected correlation between a smooth ground truth and sparse observations. Since we lack the ground truth of technical dropout-free gene expression we need to approximate it. We can use auxiliary methods to denoise the observed counts which includes imputing observed zeros. Here we denoise the HSC data using scVI[21]. The expected cross-cell correlation using the denoised observations as ground truth is 0.209 (Supplementary Figure 1 B), which we use as a sparse evaluation baseline (Figure 3 C). We termed the gap between our current model performance and the sparse evaluation baseline the model gap. It estimates the performance a hypothetical perfect model can achieve on this data set using the same evaluation. We can also compare our smooth model predictions directly against the denoised observations (Figure 3 C & D), where our model achieves 0.285 cross-cell and 0.762 cross-gene correlation. We named the gap between the sparse evaluation gap and the performance compared to denoised data the sparsity gap, since it is determined by the technical dropouts of a single-cell datase, rather than the model performance and cannot be overcome when working on raw data.

**Figure 4:**
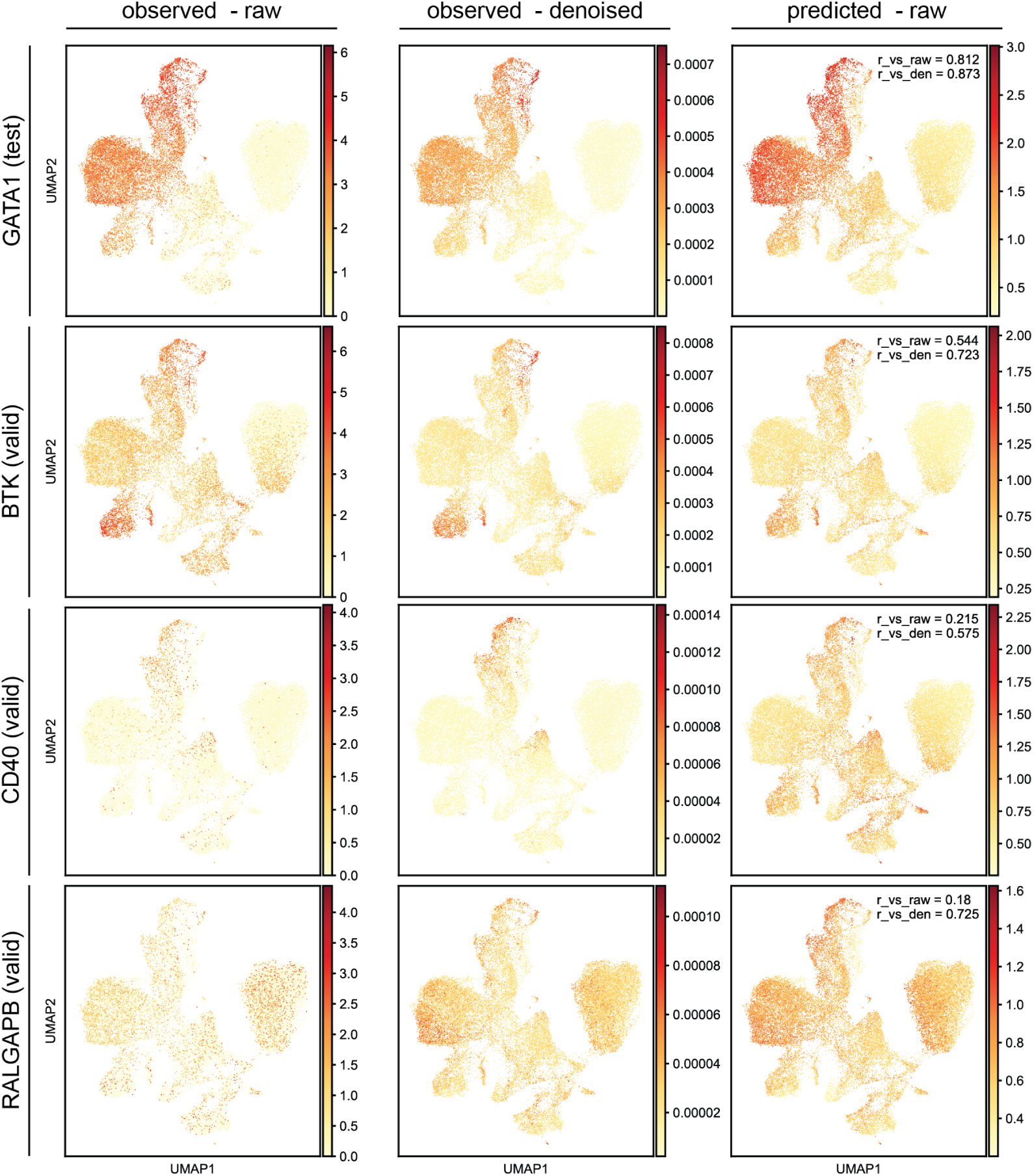
Predictive performance of example genes with varying cell type specificity from hold out sets. From top to bottom: GATA1 (test set), BTK, CD40 and RALGAPB (validation set). Shown are the raw observed (left), the denoised observed read counts (middle) and the single-cell model predicted read counts (right). Pearson correlation of the selected gene across cells of the predictions with the raw data (r_vs_raw) and the denoised data (r_vs_den) are shown in the prediction panels.

It is possible to train models directly against the denoised single-cell data (Supplementary Figure 2). Compared to denoised observations, the denoised model predictions reach cross-cell correlations at the level of pseudo-bulk analysis (Supplementary Figure 2). However, the denoised predictions do not explain the raw data better than models trained on raw observations. Furthermore, we observed artefacts of the denoising process to inflate the performance of models trained against denoised data (Supplementary Figure 3). This is the case for genes that are not expressed or are highly expressed across a single-cell dataset. Single-cell autoencoders are designed with highly variable genes in mind and are usually only applied to highly variable genes[21]. Furthermore, since the scVI autoencoder is trained on gene expression profile reconstruction from latent factors, it uses information across genes. This leads to data leakages into the hold out gene sets, compromising our ability to demonstrate generalizability. Moreover, caution is generally warranted when interpreting denoised gene expression values for downstream application[21, 22]. In conclusion, here we only apply denoising to get a ground truth approximation to compare our models trained against raw data to. Single-cell gene expression models can be trained against denoised data but we suggest caution when interpreting their results.

To compare single-cell models to the pseudo-bulk approach, we pseudo-bulked the observed single-cell counts and trained models against the average expression per pseudo-bulked cell type, mimicking the scEP[7] workflow. This yielded an across cell type correlation of 0.411 and across genes of 0.789. Alternatively, we can train a single-cell model and pseudo-bulk the model predictions. This approach yielded 0.462 across cell and 0.785 across gene correlations. Of note, in the validation set, the pseudo-bulked single-cell model performed slightly worse than the model trained on pseudo-bulked data (Supplementary Figure 2), indicating that on average the two approaches are likely on par. Interestingly, this alleviates the need to train pseudo-bulked models as predictions at the single-cell level can be summarized to any desired resolution of cell type annotation. We termed the gap between the pseudo-bulk analysis and the single-cell model predictions against denoised observations the complexity gap. It reflects the added difficulty of predicting gene expression at single-cell, rather than at pseudo-bulk level. This complexity gap is small for the cross-gene but pronounced for the cross-cell correlation. In addition, we trained a model to predict the CAGE-seq tracks of the Enformer compendium directly from the gene embeddings. On the test set this achieved a Pearson correlation of 0.488 across cell types and 0.808 across genes. In contrast, Sasse et al.[16] achieved only an across gene correlation of 0.580 between Enformer CAGE-seq predictions and population average mRNA levels of matched tissues. We indicate these as the Enformer CAGE-seq baseline and the Enformer population average baseline respectively (Figure 3 C & D).

Single-cell biologists usually assess the quality of cell clustering and annotation by the distribution of marker genes, genes specific for a subset of cell types. Similarly, we can show the performance of the single-cell model on predicting cell-specific gene expression on hold out genes not seen during training (Figure 4), demonstrating generalizability and cell specificity. As shown in previous work[5] our models cell-specific performance varies depending on the gene, as a model will inevitably learn the sequence determinants of some genes better than others. However, rather than just reporting an average metric and glossing over poorly performing genes, we can leverage the cross-cell correlation per gene as a readily available confidence metric. We are more likely to trust variant effect predictions on genes whose cell-specific expression profiles were learned well by the model. We do not consider genes with a cross-cell correlation below 0.1 as having captured cell-specific gene expression (Supplementary Figure 4). Following the logic of scBasset[23], developed for scATAC-seq, we can further evaluate models by applying dimensionality reduction to the weights of the final linear layers which can be treated as cell embeddings (Supplementary Figure 5).

### Variant effect prediction in single cells reveals heterogeneity within cell types

Discovering and interpreting the cell-type-specific effects of non-coding sequence variation on gene expression is still a key challenge for understanding disease-linked genetic variability. The most direct link between sequence variation and gene expression are expression quantitative trait loci (eQTLs). eQTL studies are traditionally performed on bulk RNA-seq data, often on broad tissues such as whole blood. This renders it difficult to discover variants with cell-type-specific effects [24] unless an affected cell type dominates the bulk data used. Even when utilizing single-cell data, pseudo-bulking is usually employed[18], although single-cell resolution eQTL mapping techniques such as variance eQTLs or eQTLs along trajectories are gaining popularity[25]. As variant effects may be highly specific to individual cell types as well as developmental and activation stages, enabling discovery at high resolution is crucial. Pseudo-bulking will miss variant effects that vary within aggregated cell types. Potentially, this greatly impacts an eQTL study’s ability to identify variants with cell-type-specific effects and especially variants with opposing effects in the same pseudo-bulked cell type. Sequence-to-expression prediction at single-cell resolutions offers an opportunity for characterizing the cell specificity of sequence variants in single-cell systems. Predicting the effect of sequence variants on cell-specific gene expression, we expect to find varying levels of cell- and cell-type-specific effects (Supplementary Table 1). Because we observe gene expression patterns at a granularity beyond annotated cell types (Figure 1), we expect to find variants effects on gene expression at the same resolution (Figure 5).

**Figure 5:**
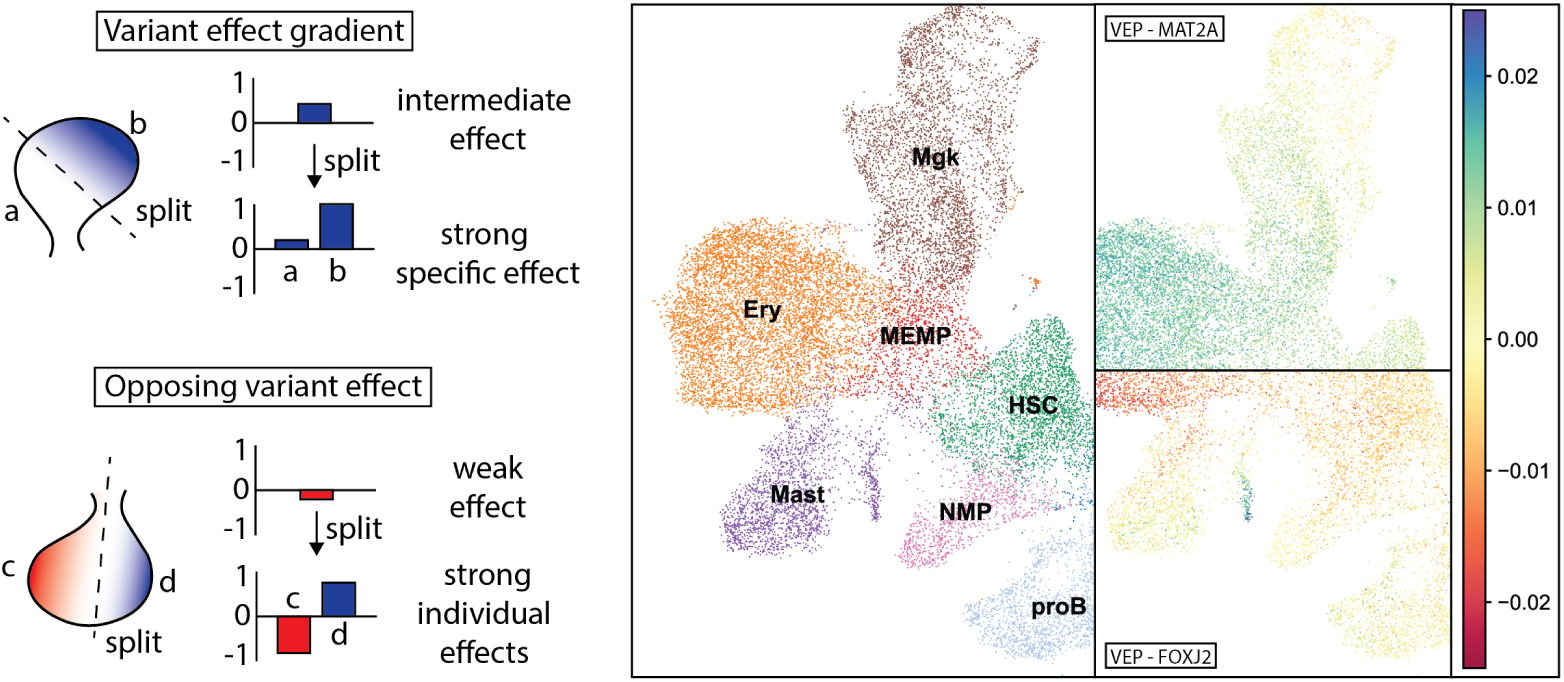
Cartoon (left) outlining potential sequence variant effects on gene expression where sub-cell-population structure may lead to dampening the average effect in pseudo-bulked data. Examples (right) of predicted variant effects of whole blood eQTLs in the single-cell HSC model highlight sub-cell-population effects that might be missed in pseudo-bulked analysis. The color gradient indicates the difference (delta = variant - reference expression) in the variant effect prediction (VEP) associated with a single base pair variant.

To demonstrate how sequence-based variant effect prediction can characterize cell-specific effects, we predicted the impact of fine-mapped, whole blood eQTLs[26, 27] with our HSC model. The large sequence context allowed us to evaluate 1,154 variants up to 100 kb away from the TSS. We extracted the ~ 200 kb centered on the canonical TSS of the eQTL linked gene and fed the one-hot-encoded sequence with the reference and the variant allele through the seq2cell model. We then calculated the difference between the variant and reference predicted expression.

First, we investigated how well the average predicted effect across all cells explains the eQTL effect size, i.e the slope of the linear regression (Figure 6 A). Strikingly, the majority of eQTLs do not show a clear predicted average effect in our HSC model. The predictions suggests that the average effect across cells cannot explain about 80 % of eQTLs. This result may be explained by the model failing to generalize to sequence variants or a tissue mismatch, where the variant affected genes (eGenes) may not be expressed in the cell types investigated or are differently regulated in embryonic vs. adult and whole blood vs. HSC differentiation. Moreover, the eQTL fine-mapping may be imperfect or eQTLs may not be caused by a cis-regulatory effect. Lastly, the effect might be more subtle than an average effect across all cells. Previous work has raised concerns about sign mismatches between Enformer-based variant effect predictions and observed differences in gene expression[15, 16]. We thus quantified the percentage of eQTLs in each quadrant of the delta vs. eQTL slope plot. We found that the majority, but not all, predictions of strong average predicted variant effects agreed with the eQTL slope sign. Previous work also raised concerns that Enformer-based gene expression predictions overly focus on sequence determinants close to the TSS, down weighting distal enhancers[14]. Plotting the mean absolute value of the delta vs. the distance to the linked TSS, we observed a trend for variants to have a higher average predicted effect closer to, and particularly in the range of the promoter (~ 2 kb) (Supplementary figure 6). Here, the prediction matches the expectation that more distal gene regulation mediated by enhancers has cell-type-specific effects[28], whereas variants in promoters tend to affect gene expression more broadly. Interestingly, we still found comparatively strong average predicted effects beyond 10 kb, the current sequence context of other single-cell or pseudo-bulk expression models[7, 8].

To check whether the average effect prediction across cells statistically enriches for functional variants, i.e. variants with observable effects on gene expression, we sampled negative SNP sets matching the eQTL dataset. For each eQTL we sampled 3 x 10 variants at distances between 1 to 100; 100 to 1,000 and 1,000 to 10,000 bp into three distinct negative sets. We assume that: 1) Randomly sampled variants would yield, on average, non-functional variants. 2) With increasing distance from the eQTL we are less likely to disrupt the same regulatory element as an eQTL might. We predicted the effect of the negative SNP sets with the HSC model and plotted the cumulative probability of the average absolute effect across cells (Figure 6 B). Sampling at increasing distances from eQTLs depletes the predicted average effects. eQTLs have significantly higher predicted effects on gene expression compared to each negative SNP set (two sample Kolmogorov–Smirnov test p-value: < 1e-17; < 1e-105; < 1e-208 for 1 to 100 bp; 100 to 1,000 bp; 1,000 - 10,000 bp respectively). We fixed 0.005 of the mean absolute delta across cells as an empirical threshold where 22 % of eQTLs pass, 14 % of negative SNPs within 100 bp pass, those are more likely to disrupt the same element as the matched eQTL; only 5 % of negative SNPs between 100 bp and 1 kb pass and only 1 % of negative SNPs between 1 and 10 kb pass.

**Figure 6:**
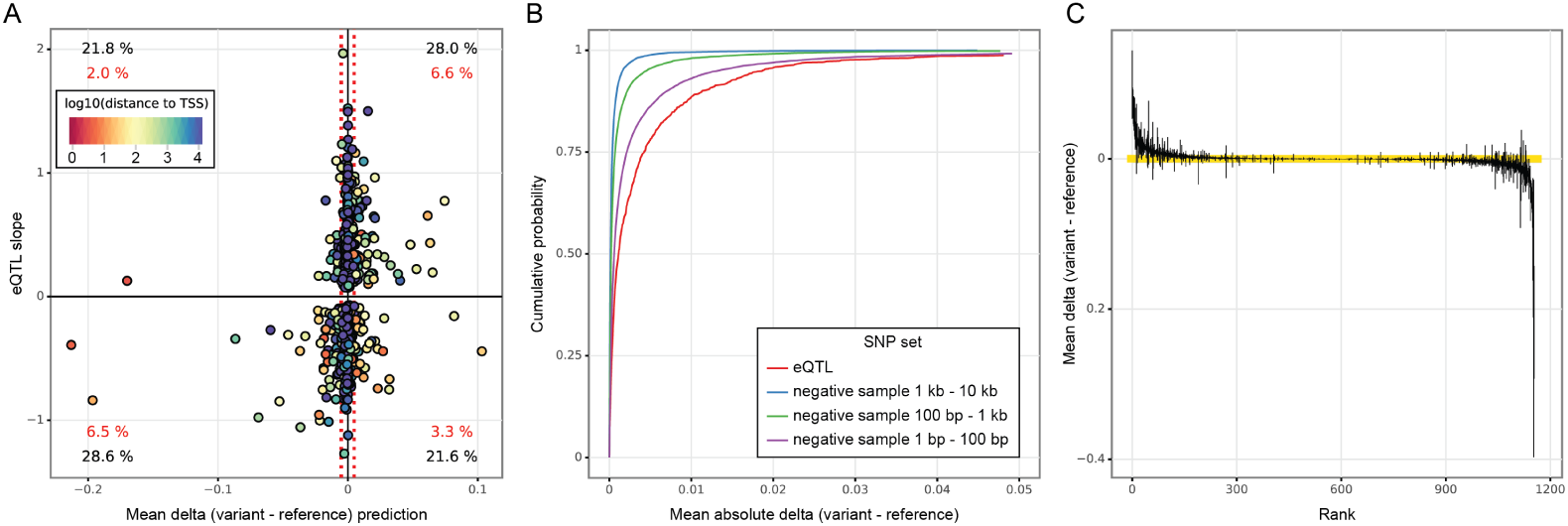
Predicted effect on expression vs. eQTL effect. **A)** 1,048 fine mapped eQTLs matching to expressed genes in the HSC dataset and whose eQTL slope could be extracted. Plotted on the y-axis is the eQTL effect size, the slope of the linear regression fitting the read counts from homozygous (hom) reference to heterozygous to hom. alternative allele. Plotted on the x-axis is the predicted effect of the variant on the eGene expression averaged across all cells (mean variant expression – mean reference expression). Each dot represents a single variant – gene pair and is colored by the distance to the TSS of the respective eGene. Red dotted lines indicate the positive and negative empirical thresholds on the average predicted effect determined by SNP permutations. Percentages quantify variants within each quadrant: black - all variants; red - all variants passing the empirical threshold. **B)** Cumulative probability of the mean absolute predicted change in expression for the eQTL set and for three sets of matched negative SNP samples at increasing distances: 1 – 100 bp; 100 – 1000 bp; 1000 – 10,000 bp. **C)** Predicted effect of 1,154 whole blood eQTLs on their eGenes in the HSC dataset. Each variant’s effect (delta = variant – reference expression) was predicted in single cells and averaged per annotated cell type. Plotted is one line per variant indicating the range of the predicted average effects across cell types (min to max). Lines crossing 0 indicate different directions of predicted variant effect depending on the cell type. Variants were ranked by the mean of the delta. Yellow bar indicates the range *±* 0.005 delta, our empirical threshold for variant effects.

Because the average predicted effect is a suboptimal eQTL effect predictor, we next sought to estimate the variability in cell-type-specific effects. To this end, we averaged the predicted single-cell effects per annotated cell type. We identified the cell type with the highest and lowest average predicted effect per variant, and plotted the range on the average-effect-ranked variants (Figure 6 C). Out of the 1,154 variants, 500 (43 %) flip between a positive and negative predicted average effect depending on cell type. For 163 (14 %) variants, the range between cell-type-specific average effects was larger than 0.005, which is our empirical threshold for averaged variant effects.

Lastly, we sought to estimate variant effect heterogeneity within annotated cell types, beyond the resolution of pseudo-bulks. To this end, we calculated the standard deviation of the predicted effect of each variant within each annotated cell type. We found 202 variants (17.5 %) have at least one intra-cell-type predicted standard deviation larger than 0.005. We visualized the top six examples of heterogenous, intra-cell-type variant effects (Supplementary Figure 7). We observe predicted variant effect gradients (COPB1, C12ord75, ARHGEF18, GTF3C3) and bivalent variant effects within the same annotated cell type (EFHD2, ACKR1). We expect such variants to be particularly hard to identify even when using single-cell data to create pseudo-bulks.

To gauge if variant effects beyond the resolution of pseudo-bulked, annotated cell types are driven by true biological differences we characterized the predicted variant effects of exemplary eQTLs in more detail. The variant 12_8058041_A_G (rs2072449) is an eQTL for the transcription factor FOXJ2. The variant is located at a distance of *~* 25 kb to the canonical TSS of FOXJ2, situated in the 3’ UTR of C3AR1. It has been associated to changes in expression of C3AR1 and to act in *cis* on FOXJ2 and NECAP1 (GTEx Portal[26]). The variant is in an ENCODE candidate cis-regulatory element[29]. The G allele was found to have an expression enhancing effect on FOXJ2 in the whole blood eQTL study. However, multi-tissue eQTL comparisons point to expression reducing effects on FOXJ2 in EBV-transformed lymphocytes and on NECAP1 in multiple tissues (GTEx Portal). In GWAS studies, the G allele of rs2072449[30] was found to be associated with a decrease in monocyte count and an increase in neutrophil count.

Our single-cell variant effect prediction matches the bivalent nature of 12_8058041_A_G (Figure 7). The difference in predicted variant effect is not explained by differences in FOXJ2 expression (Supplementary Figure 8). The variant is predicted to reduce FOXJ2 expression predominantly in HSC, megakaryocyte erythrocyte mast cell progenitors (MEMPs), erythroid cells (Ery) and double negative T cell progenitors (T_DN). In Megakaryocytes (Mgk) the variant effect is predicted to be negative, closer to the progenitor cells in UMAP space and weakens and flips the effect direction towards the extremity of the Mgk cluster. Splitting the Mgk cluster based on variant effect strengthens the negative effect in the earlier population and highlights the average positive effect in later stage cells. The later Mgk cells show a higher average expression of PPBP and other marker genes of platelets and mature Megakaryocyes. The predicted variant effect also weakens towards the extremities of the Mast cell cluster. Interestingly, the Park et al. annotated Mast cell cluster shows substructure, and the variant is predicted to have a stronger positive effect in one of the subclusters. Splitting the annotated Mast cells based on the substructure and difference in predicted variant effect, highlights the effect pattern. Relative to the cleaner set of Mast cells, the split cells are depleted for KIT expression, a Mast cell marker, and enriched for CLC expression a marker gene for Eosinophil and Neutrophil cells. This suggests that the original annotation by Park et al. was not of sufficient resolution.

**Figure 7:**
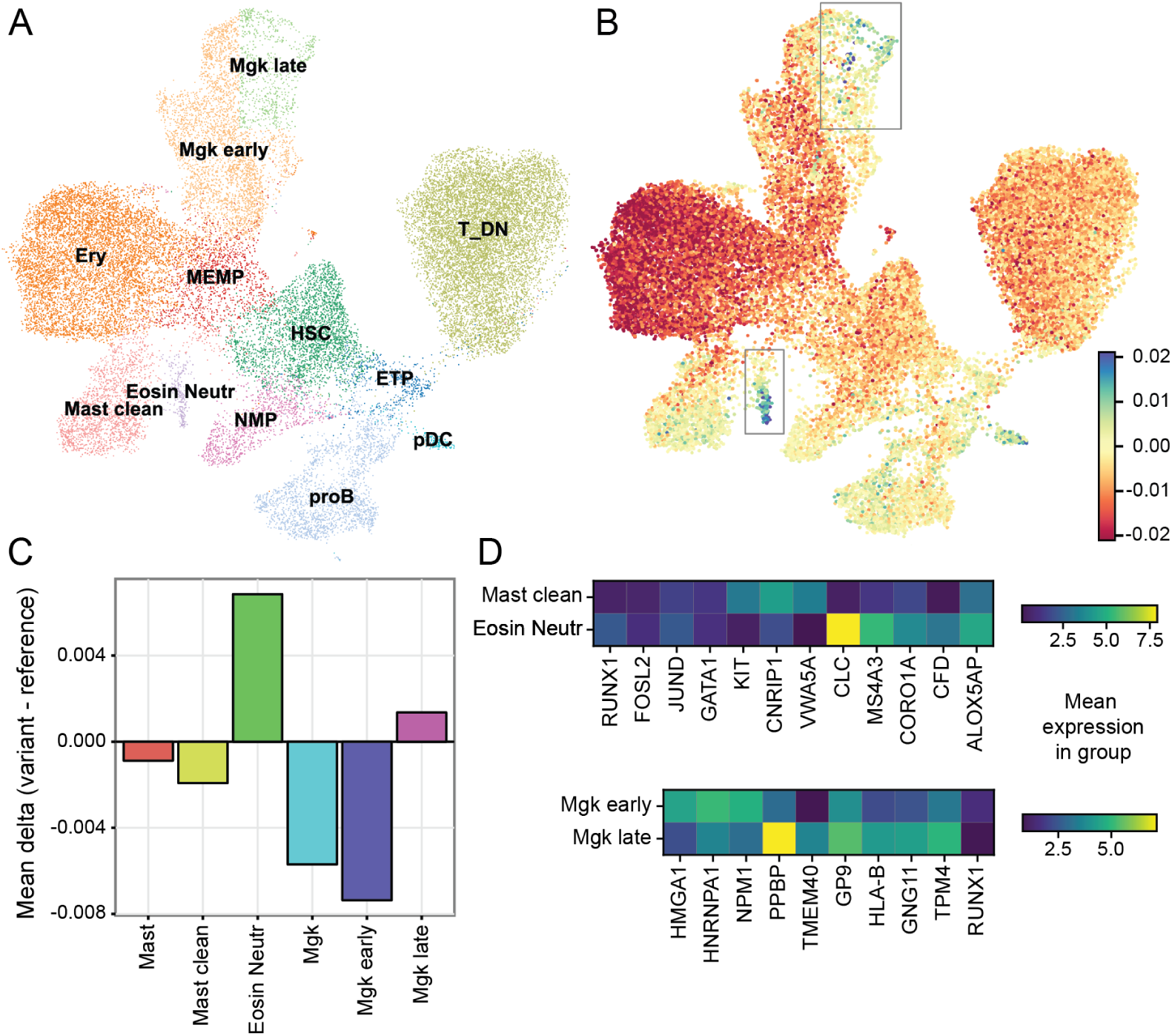
Heterogeneity within annotated cell types of the predicted effect of the FOXJ2 linked whole blood eQTL: 12_8058041_A_G. **A)** UMAP after splitting the cell type annotations for Mast cells and Megakaryocytes based on **B)** the predicted effect of the eQTL on FOXJ2 expression, indicated by the color gradient. **C)** Mean predicted affects in the split subpopulations (Mast cells to Mast cells and Eosinophils/Neutrophils; Megakaryocytes (Mgk) into early and late) are in a different direction and drowned out in the cell population pre-splitting. **D)** Characterizing the split populations by marker genes indicates (left) that the cell population split from the clean Mast cells is depleted for KIT, a Mast cell marker and enriched for CLC, a marker of Eosinophils and Neutrophils. (Right) Compared to the early Mgk cells, the late Mgk are enriched for PPBP and other markers genes of mature Megakaryocytes.

On the sequence level, the G allele weakens an ETS-domain family binding motif. The ETS-domain family of transcription factors comprises a multitude of factors with a conserved binding motif. ETS-domain family TFs may act as transcriptional activators or repressors depending on the TF and its interactions with co-regulators[31]. A multifaceted effect based on expression of ETS-domain family members (Supplementary Figure 9) and co-regulators may explain the bivalent variant effect. In addition, 12_8058041_A_G disrupts weak GATA1 and GATA3 motifs, which may explain the negative predicted effect on expression of FOXJ2 in HSC, Erythroid and T cell progenitors (Supplementary Figure 10).

Taken together, 12_8058041_A_G, provides an important example for multifaceted variant effects that may lead to discrepancies between eQTL slopes and sequence model predictions. Bivalent effects through weakening of ETS-domain TF binding as well as disruption of a GATA binding site are convincing indicators of a highly context dependent effect on gene expression that can only be fully characterized through single-cell resolution.

As a second example, 2_85501572_A_T (rs72844419) an eQTL for MAT2A, is characterized by a similar sub-cell-type effect but in opposite direction (Supplementary Figure 11-13).

### Scaling seq2cells maps variant effects from T cell activation to T cell development

We next sought to apply seq2cells to more challenging datasets, both in terms of scale, i.e. the number of cells, and homogeneity in cell-to-cell differences (Supplementary Figure 14). We trained seq2cell models against the full 250 k cell dataset of the human T cell development atlas [12] and against a 650 k cell dataset following the activation of CD4 T cells, memory and naïve over time[18]. Owing to the flat emb2cell architecture we chose, it was necessary to reduce the bottleneck dimension to fit the 650 k cell model (see Methods). Despite more challenging datasets, seq2cell was capable of capturing cell specific gene expression. Overall, the development model achieved a test set mean Pearson correlation across cells of 0.169 and across genes of 0.533. The correlations were 0.078 and 0.666 in the activation dataset respectively. Of note, the activation dataset follows two related cell types over a time course past stimulation, yielding a much more homogenous dataset with less distinct cell types clearly separable in UMAP space (Supplementary Figure 14). We expect less cell-to-cell variation in the gene expression profiles of closely related T cell states, compared to the HSC and T cell development datasets with more diverse cell types. In the activation data, the variation in gene expression is driven by more subtle differences, which are harder to capture and explain a drop in the average cross-cell correlation. We observe far fewer genes with a clear cell type specificity compared to more heterogenous datasets. However, we can use the per gene Pearson correlation across cells as a metric to indicate for which genes the cell-specific expression patterns have been captured reasonably well, bolstering confidence in variant effect predictions. Using a cross-cell correlation of 0.1 as threshold: In the 250 k T cell development data, 9,624 (64 %) of training, 862 (59 %) of validation and 1083 (61 %) of test set genes pass the threshold. In the 650 k T cell activation dataset, 2,455 (15 %) of training, 307 (24 %) of validation and 399 (24 %) of test set genes pass the threshold.

To test whether seq2cells is capable of variant effect predictions in these more challenging settings, we utilized the pseudo-bulk-based eQTLs identified by Soskic et al.[18] using the genotypes of the 119 individuals from which the CD4 T cells were derived. In total, Soskic et al. identified over 480 k significant variant gene interactions. Because, those eQTLs have not been fine mapped, we can only attempt to surface eQTLs that disrupt learned sequence determinants of gene expression. We focused on associations with genes that passed 0.1 cross-cell correlation and variants within 100 kb of their eGene’s TSS, excluding indels. To further reduce the number of variants and avoid potential spurious correlations from overfitting, we restricted to validation and test set genes, yielding a total of 5,930 variant – eGene associations covering 502 individual genes and 4,711 individual variants. We predicted the variant effect on gene expression at single-cell resolution using the T cell activation and T cell development models. First, we used the coarse measure of average predicted effects within each whole dataset and within pseudo-bulked cell types. With the activation model, we identified 30 associations (0.5 %) whose average predicted effect across all cells pass our empirical threshold including 28 unique variants and 27 unique eGenes (Supplementary Figure 15). Translating the variant effects to the development model, we identified 54 associations (1 %) whose average effect across all cells pass our threshold, including 52 unique variants and 45 unique eGenes. The correlation of the average prediction across cells, between the development and activation model predictions is 0.56 (Supplementary Figure 16 A).

We then compared the pseudo-bulked predictions of the 41 associations where at least one pseudo-bulked cell type effect passes our empirical delta threshold in one of the datasets (Figure 8). The results suggest that T cell activation eQTLs are also affecting developing T cells. We observe associations with the same and opposing effects in activation and development. Variant effects tend to be accentuated in T cell activation and early T cell development (early thymic progenitors (ETP) to proliferating double positive (DP) thymocytes). If opposing variant effect directions are predicted, they deviate in early T cell progenitors and in activated T cells, primarily 16 h and 5 days after stimulation. We correlated the predicted affect across the 41 associations between cell types in both datasets (Supplementary Figure 16 B). We observe higher correlation within each dataset. Within the datasets, the variant effects group individual time points of T cell activation as well as the early T cell progenitors (ETP, gamma delta T cells and proliferating double negative (DN) and DP thymocytes) and the later stages of T cell maturation. Interestingly, quiescent DN and DP thymocytes have a variant effect prediction profile more closely related to the later stages of T cell development as opposed to their proliferating counterparts and earlier progenitors. Furthermore, the predicted variant effects in early T cell progenitors show a higher correlation with the predicted variant effects in activated T cells, as opposed to non-activated T cells. This suggests that similar transcriptional programs may be involved in T cell activation and development.

**Figure 8:**
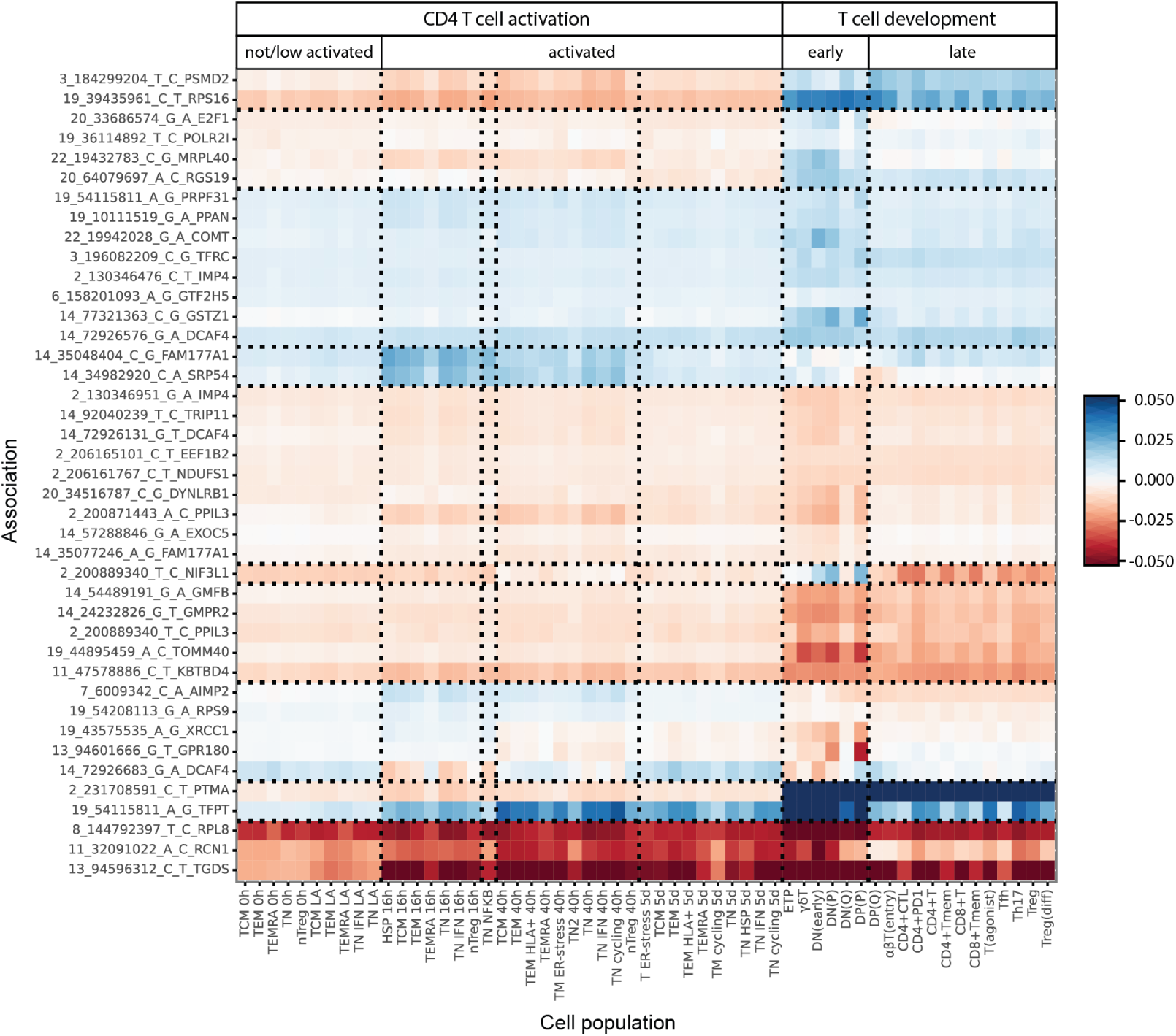
Comparative variant effect prediction in CD4 T cell activation and T cell development. Color indicates the predicted mean delta (variant – reference expression) of pseudo-bulked single-cell predictions. Shown are 41 variant – eGene associations, of validation and test set genes that were learned sufficiently well by the models (Pearson Correlation across cells > 0.1) and which pass the empirical delta threshold of *±* 0.005 in at least one pseudo-bulked cell type.

Next, we sought to characterize individual variants across T cell development and activation at single-cell resolution. We identified variants, where the predicted effect follows cell type delineation (Supplementary Figure 17). The variant 2_200889340_T_C (rs7559150) was identified by Soskic et al. to reduce NIF3L1 expression in memory T cells, 16 h after stimulation. The variant lies within a dual promoter of NIF3L1 (31 bp to TSS) and PPIL3 (36 bp to TSS) and eQTL studies have shown effects on both genes (GTEx Portal). In the activation model, the variant is predicted to have a negative effect on expression in CD4 T cell activation, with the effect being stronger in non-activated T cells, where NIF3L1 is also expressed less. In T cell development, the variant has a predicted bivalent effect depending on the cell type. The variant has an enhancing effect on expression in proliferating DN and DP thymocytes, but a reducing effect in quiescent DN and DP cells and in more mature T cells.

Other variants showed predicted effects more granular than the annotated cell types (Supplementary Figure 18). The variant 14_72926683_G_A was identified by Soskic et al. to have a positive impact on DCAF4 expression in a subset of naïve T cells, 40 h after stimulation. The intronic variant is 220 bp downstream of the DCAF4 TSS. Interestingly, the predicted effect is negative in some T cell populations 16 h after stimulation. In the development dataset, the predicted effect is negative in most non-T cell types, but positive further along the T cell trajectory. The variant changes an E-box motif from CACGTG to CACATG (Supplementary Figure 19) implicating E-box binding factors in its mechanism of action. We therefore looked at the expression MYC, MAX and MXD1 (Mad1) along the T cell development trajectory. MYC is expressed predominantly in early T cell progenitors and non-T cell types in the development dataset. A weakening of the E-box matches with a predicted decrease in DCAF4 expression in those cells. MAX can bind the DNA as homodimer or dimerize with MYC or MXD1. The reduction of MYC expression along the T cell development trajectory coupled with an increase in MAX and MXD1 expression tracks the different predicted variant effects. This suggests that MXD1-Max heterodimers take over from Myc-Max binding as cells progress through the T cell development trajectory. MXD1 is a dedicated transcriptional repressor with identical binding preferences to MYC, a transcriptional activator[32]. Therefore, a weakening of the E-box motif matches with an increase in DCAF4 expression. A similar change in predicted effect tracks with MYC and MXD1 expression in the activation dataset (Supplementary Figure 20).

A large sequence context enables us to interpret variants at larger distances from their associated eGenes. The variant 2_200871443_A_C (rs7562391) is 17,635 bp away from the TSS of PPIL3, on which Soskic et al. identified a negative impact on expression in the T cell population characterized by heat shock proteins, 16 h after stimulation. The variant is located within the last exon of PPIL3 and predicted to be a misssense variant. However, the region is also marked by H3K4me1 in K562 cells and classified as a distal enhancer in ENCODE candidate regulatory element[29]. Enhancers co-located with exons have been described [33]. The variant is predicted to have a negative effect on PPIL3 expression, predominantly in activated T cells and in early stages of the T cell development trajectory (Supplementary Figure 21). The variant weakens a potential HOX binding motif (Supplementary Figure 22). Among those HOXA9 has been implicated for T cell development [34] and various HOX genes are expressed along the T cell development trajectory. In addition, the variant also improves a FOXP1 binding motif. FOXP1, a transcriptional repressor, is expressed in T cell development but depleted in the T cell progenitors where the variant has its strongest effect. These results hint at a multifaceted variant effect involving FOXP1 and HOX TFs.

Taken together, by scaling seq2cells, we can predict the variant effect of CD4 T cell activation eQTLs in the context of T cell development. Seq2cells predicts variant effects at resolutions beyond annotated cell clusters and predicted effects can be highly cell-specific and flip their sign based on the context. This highlights the value of single-cell level variant effect predictions for cell annotation and variant interpretation.

## Disussion

In this work, we present seq2cells as a framework for granular gene expression prediction from large scale DNA sequence inputs at single-cell resolution. Previous machine learning methods for predicting single-cell gene expression from DNA sequence had to compromise on a combination of resolution, granularity and sequence context [7, 8]. scEP[7] tries to strike a balance between the resolution of single-cell data and its sparsity problems. The approach uses pseudo-bulked single-cell data to train a convolutional neural network (CNN) to predict pseudo-bulk gene expression. Pseudo-bulked data may offer robustness but sacrifices resolution. Yet, single-cell data readily reveals examples of sub cell type structure in gene expression that pseudo-bulking is unable to capture. Marker genes are not simply tuned to specific expression rates per cell type but can show variability within (Figure 1). Furthermore, pseudo-bulking single-cell data abolishes the ability to learn from and predict on a continuum of cells along trajectories. Nvwa[8] trains a CNN to predict gene expression at single-cell resolution. To deal with the data sparsity Nvwa employs imputation of missing data, yet imputation methods can produce artifacts (Supplementary Figure 3) and negatively impact downstream analysis[35]. In addition, Nvwa requires discretization of gene expression into on or off, limiting its ability to interpret the potentially weak effects of disease-linked variants [36] on a continuum of gene expression. Both scEP and Nvwa operate with a limited sequence context. Nvwa uses a 10 kb context centered on the TSS. scEP uses a 10.5 kb context reaching 7 kb upstream of the TSS. Yet we know that cell-type-specific regulation happens at longer sequence ranges[37] and disease associated variants are enriched in regulatory elements[38].

Seq2cells rests on the premise that, using pretrained epigenome models to create DNA sequence embeddings of the TSS, we can create gene embeddings that encapsulate transcriptional regulation. This allows us to predict gene expression from the sequence determinants, for example TF motifs, that drive gene expression mechanistically, rather than from co-expression patterns of genes[39–41]. In the future, we hope to aid other gene-level task with the information contained in the regulatory DNA sequence of a gene. The seq2cells framework can in principle use any model that embeds the DNA sequence of the TSS as its seq2emb module. Here, we adopted Enformer as our epigenomic base model. Apart from the single-cell aspect, compared to Enformer we shifted our attention to tackling the cross-cell (type) correlation which received little attention in the original work[5]. Moreover, because we want to make variant effect predictions, avoiding over-fitting is crucial. We want to make predictions on training set genes without elaborate cross-validation schemes at inference time.

Furthermore, we want to avoid the effect of spurious attributions[16] which are likely the effect of overfitting. This is noteworthy compared to recent work: For example, scBasset[23] predicts single-cell accessibility from DNA sequence of *~* 1 kb. scBasset was designed to optimize the low dimensional latent space embedding and accepts overfitting on the training set, if training set performance continues to improve. Similarly, training a single linear layer from Enformer TSS embeddings to CAGE-seq data without additional regularization markedly overfits to the training set. We employ weight decay and a high dropout rate (0.5) to ensure that training and validation correlations are on par.

We chose Enformer for its connection to gene expression via the CAGE-seq tracks and its longer sequence context of *~* 200 kb, compared to most existing genomic language models[9, 10] Recently, HyenaDNA[42] has been proposed as DNA language model with up to one million base pairs of sequence context. This model is promising but it is yet unclear if transfer learning to single-cell expression will work out of the box or if prior fine tuning to epigenomic assays such as CAGE-seq or RNA-seq is required. Another potential limitation is the pre-training corpus of Enformer, which focuses on broad cell types and cell lines. Here, we transfer learn on new data, but seq2cell models may perform better the closer the single cells are related to cell types covered in the Enformer corpus. Finetuning the entire Enformer model, rather than keeping the trunk frozen may address this better at the cost of substantially larger computation costs.

In this work, we have taken a two-layer MLP approach for the emb2cells module. Using an individual output per cell in a large, final linear layer scales linearly with the number of cells but with a factor equal to the bottleneck dimension, i.e. every cell adds 2,000 weights and biases to the model. Given the sizes of modern cell atlas datasets of over 1 M cells per dataset, scaling this architecture will be limited. Here we successfully trained a single-cell expression model on 250 k cells and with compromising on the bottleneck dimension on 650 k cells. Purely transcriptomics-based single-cell models [39–41] demonstrated the benefits of scaling to millions of cells. We envisage similar benefits from scaling sequence-based models into this realm. However, this will require an architecture scalable to tens of millions of cells.

We evaluated model performance against raw observed, denoised observed and pseudo-bulked observed data. We developed a sparsity evaluation baseline to estimate how well a perfect model could predict a given, sparse, single-cell dataset. This allowed us to structure the gap between our single-cell model performance and pseudo-bulked or bulk level models: 1) The model gap describes the space for improvement between our current model performance and a hypothetical, perfect model given our sparse dataset. 2) The sparsity gap is intrinsic to evaluating single-cell predictions against sparse observations. 3) The complexity gap describes the added difficulty of predicting single-cell gene expression rather than pseudo-bulked aggregates. For improvements in future models, the gaps could be addressed individually. Narrowing the model gap by better architectures will improve all model evaluation. The sparsity gap may be overcome by more efficient capturing technology or by using denoised observations. Indeed, when we train seq2cell models against denoised data the performance metrics markedly increase. Single-cell imputation requires auxiliary machine learning methods. From this process we will inevitably inherit errors and artefacts. We observed striking artefacts of the imputation process (Supplementary Figure 3) specifically at genes with low inter-cell variability. These artefacts inflate the denoised model performance metrics. We therefore restrict to training and evaluating on raw observed data and acknowledging the sparsity gap thus induced. It is straight forward to use the seq2cells framework with denoised observations, but we caution adopters to be aware of artefacts and inflated performance metrics.

Variant effect predictions are one of the chief motivations for DNA sequence-based gene expression models. Here, we introduce a method to predict the effect of sequence variants on gene expression at single-cell resolution. Variant effect predictions so far have focused on bulk data of broad cell types. We demonstrated that single-cell predictions can be flexibly combined into cell types or whole dataset aggregates. This alleviates the need for defining cell types *a priori* to model training. Concerns have been raised how well Enformer, the current best in class model, can predict gene expression from personal genomes[15, 16]. Sign mismatches between Enformer predictions and true variant effects on gene expression suggest that Enfomer may be more suited for predicting the magnitude rather than the direction of a change in gene expression[15, 16]. When working with variant effect aggregates across whole datasets we also observe sign mismatches between eQTL effect sizes and seq2cell predicted variant effects. However, seq2cell allows us to characterize the variant effects at greater resolution. Aggregating variant effects within individual cell types demonstrates variant effects that flip direction depending on the cell context. Characterizing variant effects at single-cell resolution suggests that variant effects can be heterogenous even within an annotated cell type. Here, we used fine-grained interpretation of bivalent variant effects that can be explained by alterations to DNA binding motifs, matched with cell-specific expression of transcription factors. Our insights match with trends in single-cell genetics, where eQTL studies increasingly focus on single-cells to discover dynamic and cell specific variant effects[25].

Here, we demonstrated seq2cells utility for predicting variant effects and transferring variant effects to new cell contexts. We believe seq2cells will be a valuable tool for understanding the effect of genetic variation on transcriptional regulation in dynamic cell landscapes.

## Methods

### Single-cell datasets

Processed single-cell count matrices of the human thymic T cell developmental data[12] were retrieved from the published zenodo repository (DOI: 10.5281/zenodo.3572422). In brief, the Park et al. count processing included: normalization for total counts per cell; log (x+1) transformation and batch correction by regressing out the covariates ‘method’, ‘source’, ‘donor’ and ‘Cycle_score’. The author-provided cell annotations were used at the highest resolution available (Ann_level_5). The HSC subset and the full (ALL) set of cells were used for separate models. The raw single-cell count matrix of the CD4 T cell activation data[18] was downloaded from https://trynkalab.sanger.ac.uk. Following the preprocessing in Soskic et al., genes occurring in fewer than 10 cells were dropped, the raw counts were normalized for library size and log(x+1) transformed. The author’s cell annotations have been adopted. All time points were used for training a single model. Single-cell data and predictions were processed using scanpy[43].

### Enformer implementation and pre-trained weights

The public PyTorch implementation of Enformer (https://github.com/lucidrains/enformer-pytorch) was adapted as the seq2emb module. The PyTorch ported, published weights[5] were used (https://huggingface.co/EleutherAI/enformer-official-rough).

### cInputs and Targets

Input is a one-hot encoded DNA sequence of length 196,608 bp centered on the TSS of a given gene. Where the TSS cannot be centered without padding the sequence, the sequence window is shifted to maximize the amount of non-padded DNA sequence while keeping the TSS as central as possible. The canonical TSS of all protein coding genes (Gencode V41) were used yielding a total of 19,986 examples. These were split into 16,449 training, 1,607 validation and 1,930 test genes by overlap with the Enformer training, validation and test regions. All DNA sequences are based on the GRCh38 reference genome. Output is the predicted expression of the associated gene in every cell of a single-cell dataset. Every cell is an individual output. Targets are the normalized, log(x+1) transformed and batch corrected single-cell counts, published by the authors as processed data.

### Architecture

The one-hot encoded DNA sequence is processed by the seq2emb module, here the Enformer[5] trunk. The Enformer trunk takes as input a one-hot encoded 196,608 bp DNA sequence and outputs 3,072 dimensional sequence embeddings of the central 896 sequence windows of length 128 bp each. To deal with the even number of sequence windows processed by Enformer, the sequence window is shifted by 64 bp so that the TSS is centered within the 448th sequence window. The sequence embedding overlapping the TSS is extracted and treated as the gene embedding. The gene embedding is processed by the emb2exp module. The module consists of a linear layer mapping from the 3,072 dimensional embeddings to a bottleneck of 2,000 hidden dimensions, followed by Dropout, ReLU and another linear layer with Softplus activation to map to the targets. For training speed, the gene embeddings are pre-computed and only the emb2cell module is trained.

The full parameters consist of the Enformer trunk parameters (*~* 230 M) and the emb2cell module parameters: *~* 68 M HSC model; *~* 518 M T cell development model and ~ 655 M T cell activation model (Table 1). Only the emb2cell parameters are trained. For regularization, all models were trained with AdamW[44] with a weight decay of 0.1. The HSC and T cell development model were trained with a dropout rate of 0.5. For the 650 k cell model, the bottleneck dimension was reduced to 1,000 to fit into GPU memory. Dropout did not improve generalization in this model, suggesting that the bottleneck can be a sufficient regularizer[23]. However, experimentation with the HSC and T cell development data indicated that bottleneck dimensions below 2,000 were damaging performance. Therefore a 2,000 bottleneck dimension paired with dropout is recommended for new data where model size permits. A learning rate of 0.0001 was used with the PyTorch scheduler ReduceLROnPlateau with parameters: mode=‘max’, factor=0.25 and patience=2. All models were trained with a batch size of 50, shuffling the training data after every epoch. Cross-cell and cross-gene correlation was monitored during training and early stopping employed with a minimum change in cross-cell correlation of 0.001 and a patience of 5 epochs.

**Table 1:** Summary of key differences of the trained models.

|  | HSC | T cell development | T cell activation |
| --- | --- | --- | --- |
| Cells | ~ 30 k | ~ 250 k | ~ 650 k |
| Parameters emb2cell | ~ 68 M | ~ 518 M | ~ 655 M |
| Bottleneck dimension | 2,000 | 2,000 | 1,000 |
| Dropout rate | 0.5 | 0.5 | 0.0 |

For training speed, the sequence embeddings of TSS were pre-computed. Enformer sums the CAGE-seq signal over the TSS intersecting and the two adjacent sequences windows. To capture potential adjacency effects, the pre-computed embeddings were calculated as the average over the TSS intersecting and the two adjacent windows. We found that training against averaged embeddings reduces overfitting. At inference, we found the predictions to agree well with or without averaging adjacent embeddings. For variant effect predictions we only use the TSS overlapping window embedding not averaged with adjacent windows. This adds an additional layer of generalisation since the models trained against the averaged sequence embeddings have never seen the non-averaged embeddings used for variant effect predictions.

### Loss

A cosine-similarity-based loss was implemented to put equal emphasis on across cell and across genes correlation when the number of cells grows but the number of genes remains constant.

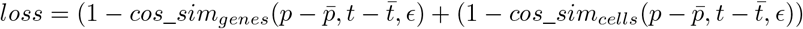

Where *p* are the predictions and *t* the targets. The average predictions 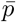 and targets 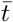 across genes and cells are estimated for each batch respectively. Epsilon is set at *ϵ* = 1e*−*8.

### Training and resources

Models were trained on up to 4 Tesla V100 with 32 GB video memory. Models took between 10 and 30 epochs to fully converge. This translates to *∼* 5.5 h for the HSC model; *∼* 9 h for the T cell development and *∼* 30 h for the T cell activation model.

### Model evaluation

For model evaluation, the Pearson correlation between the observations and the predictions across cells and across genes was calculated. For pseudo-bulking, the mean observed or predicted expression was calculated for all cells annotated as the same cell type / population.

### Imputation

The HSC single-cell data were denoised using scVI[21]. An autencoder was trained using scVI default options with the covariate keys ‘donor’, ‘organ’, ‘sort’, ‘method’ and the get_normalized_expression functionality was used to retrieve imputed and denoised counts.

### Sparsity evaluation baselines

Gene specific dropout rates were estimated as the fraction of zero entries for each gene along all cells in the HSC dataset. The denoised HSC count data were assumed as smooth ground truths for calculating the sparsity evaluation baselines. For each gene the assumed ground truth expression across cells was extracted. Then a number of cells proportional to the gene-specific dropout rate were set to 0 and the Pearson correlation of the dropout-masked expression with the assumed ground truth was calculated. This process was repeated ten times for every gene in the test and validation set, yielding an average and a per-gene estimate of the performance of a hypothetical perfect model.

### CAGE-seq baseline

A single linear layer followed by softplus was trained to predict the Enformer CAGE-seq tracks from pre-computed TSS embeddings. An ensemble of all possible Enformer inference time augmentations, up to *±* 3 bp shift and forward and reverse complement were used averaging the prediction of eight inference runs. The model was evaluated on log-transformed CAGE-seq signal.

### Weights embedding

Final layer weights of the HSC model were embedded by running PCA using the scanpy implementation with the ‘arpack’ solver and otherwise default setting; running batch aware nearest neighbor clustering bbknn[45] with the covariate ‘method’; and running the scanpy UMAP implementation with default settings.

### Variant effect prediction

For variant effect predictions the full seq2cell model was run at inference time. The reference sequence and variant sequence, centered on a TSS of interest were processed and the predicted change *delta* in gene expression per single cell is calculated as: *delta* = *p*_*var*_ *-p*_*ref*_ where *p* is the predicted expression. Average predicted effects per dataset or per annotated cell type were calculated as the mean delta and mean absolute value of the delta across cells.

### eQTL analysis

For the finemapped, whole blood eQTL dataset, all eQTLs with a significant association in whole blood were extracted from https://www.finucanelab.org/data. Matching eQTL slopes were extracted from the published eQTL study results. For the CD4 T cell activation eQTLs, significant variant – eGene associations and slopes were downloaded from https://trynkalab.sanger.ac.uk. For both eQTL sets, indels, variants further away than 100 kb from the canonical TSS of their associated eGene as well as variants effecting eGenes that did not pass the single cell quality control and were thus not presented in the respective single-cell dataset, were removed.

### SNP Enrichment analysis

Three negative SNP sets were sampled based on the filtered, whole blood eQTLs. For each eQTL 10 variants were sampled for each increasing distance window (1 to 100; 100 to 1,000; 1,000 to 10,000 bp from the eQTL variant position). The predicted variant effect was averaged and aggregated into negative SNP sets matching the allowed distance window. A two sample Kolmogorov–Smirnov test was used to test for enrichment (scipy kstest).

### Additional key packages

- anndata 0.8.0[46]
- bbknn 1.5.1[45]
- enformer-pytorch 0.5.5
- h5py 3.7.0[47]
- omegaconf 2.2.3
- patchworklib 0.5.0
- plotnine 0.8.0[48]
- pysam 0.19.0[49, 50]
- pybedtools 0.9.0[51, 52]
- python 3.8.13[53]
- pytorch 1.9.1[54]
- pytorch_lightning 1.8.1
- pandas 1.5.3[55]
- scanpy 1.9.1[43].
- scipy 1.10.0[56]

## Data availability

The published single-cell data used in this work is available as shared by the original publication. The T cell development data via the human developmental cell atlas (https://developmental.cellatlas.io/thymus-development) and zenodo (DOI: 10.5281/zenodo.3572422). The CD4 T cell activation single-cell data and eQTL results are available at (https://trynkalab.sanger.ac.uk). The fine mapped, whole blood eQTL data are available at (https://trynkalab.sanger.ac.uk). Trained models and pre-processed single-cell data for model training are available via Zenodo (https://doi.org/10.5281/zenodo.8314644).

## Code availability

Code for preprocessing, model training, evaluation and variant effect prediction is available under https://github.com/GSK-AI/seq2cells.

## Acknowledgments

We thank Lachlan Stuart, Felix Frauhammer, Frederik Träuble and the GSK.ai Sequence Learning Team for valuable discussions and support. Data used from the GTEx Portal were retrieved 14.07.2023. The Genotype-Tissue Expression (GTEx) Project was supported by the Common Fund of the Office of the Director of the National Institutes of Health, and by NCI, NHGRI, NHLBI, NIDA, NIMH, and NINDS.

