## Supplementary Material for "Single-cell gene expression prediction from DNA sequence at large contexts"

Supplementary Table 1: Expected levels of cell-(type)-specific variant effect predictions

| |
|---|
| Variants with no predicted effect, indicating that the variant does not have an effect, we are looking at the wrong cell types or states or that our model does not capture the sequence to expression relation. |
| Variants with broad and similar predicted effects across cell types. We expect those variants to be more easily identifiable in bulk and pseudo-bulk eQTL studies. |
| Variants with predicted cell-type-specific effects but in a single direction, for example reducing predicted gene expression in two out of five cell types. Such variants would be harder to discover in bulk analysis if the cell-type-specific effect does not dominate the bulk signal, for example by affecting the major cell type of the bulk. Pseudo-bulked single-cell analysis would be more appropriate for discovering such variants. |
| Variants with predicted cell-type-specific effects but opposing directions between cell types. Such variants would be hard to detect at bulk resolution since the effects would cancel each other out. Pseudo-bulking single-cell analysis would be able to handle such variants better. |
| Variants with sub-cell-type effects (Figure 5) such as effect gradients or opposing effects within the same annotated cell type . Such variants would be hard to capture with pseudo-bulked single-cell analysis. They rather require modelling at single-cell resolution and splitting of annotated cell types might be beneficial for variant effect characterization. |

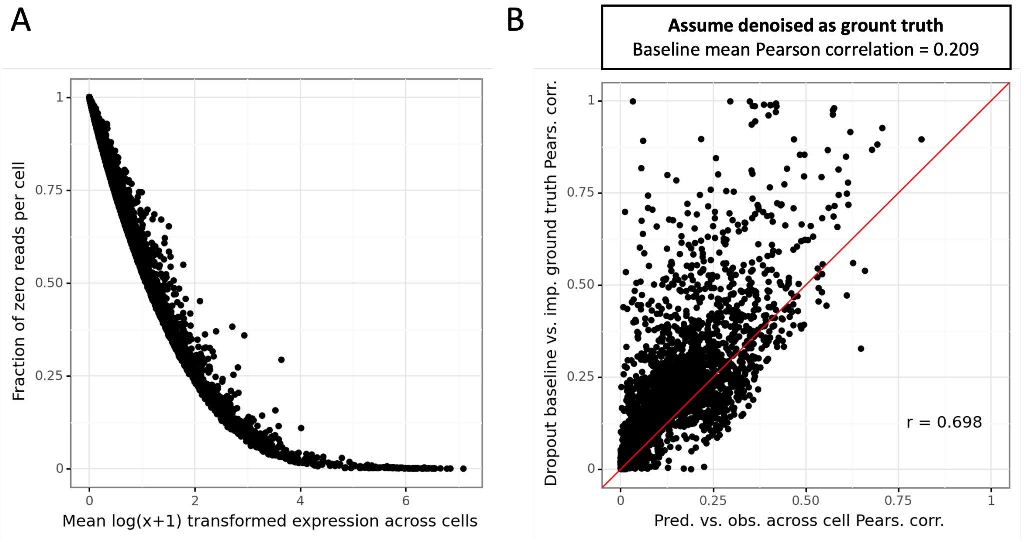


Supplementary Figure 1: Dropout rate and across cell correlation baselines for the HSC dataset. A) Stratifying the per gene dropout rate by the mean expression level. B) We assume the denoised observations as smooth ground truth. Per gene we take the smooth ground truth and set each gene to 0 with a probability equal to the estimated dropout rate and correlate the ground truth to the dropout baseline, repeating the process 10 times and calculating the average per gene. We compare this correlation to the correlation of the single cell model prediction against the raw observed counts. The mean Pearson correlation between the assumed denoised ground truth and the dropout baseline is 0.209.

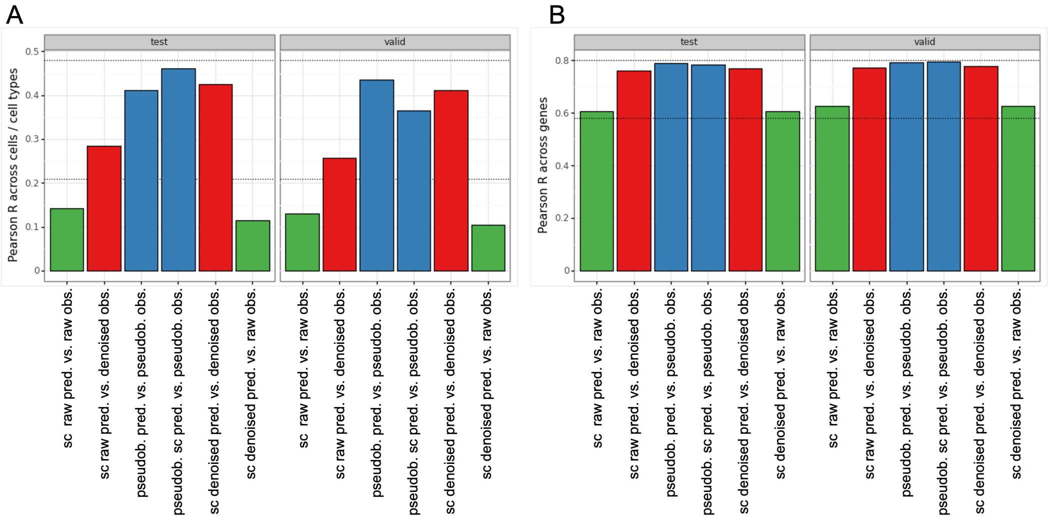


Supplementary Figure 2: Pearson correlation across cells (A) and across genes (B) stratified by validation and test set. Predictions were made with single cell models trained against raw observation (sc raw pred.) or denoised observations (sc denoised pred.) or with models trained against pseudo-bulked observations (pseudob. pred) or made with single cell models, trained against raw data, and pseudo-bulked afterwards (pseudob. sc pred.). Predictions were compared against raw single cell observed data (green), denoised single cell observed data (red) or against pseudo-bulked data (blue).

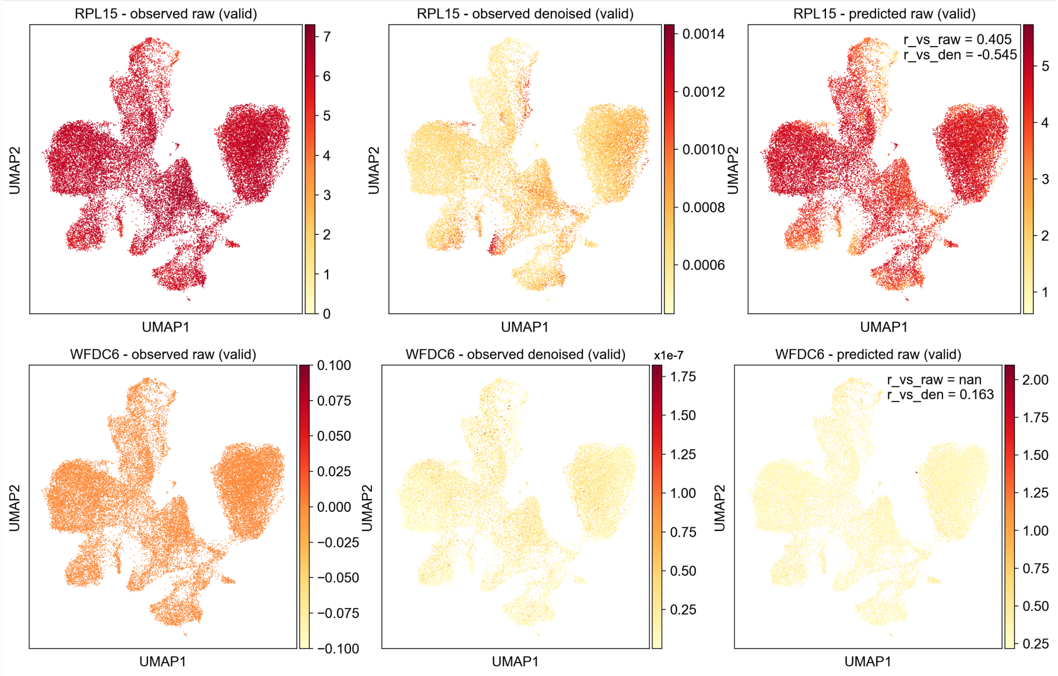


Supplementary Figure 3: Observed, raw and denoised, and predicted gene expression for RPL15 and WFDC6, highlighting imputation artefacs. Pearson correlation for the selected gene of predictions against raw observations (r_vs_raw) and against the denoised observations (r_vs_den) are shown in the prediction plot. (Top) RPL15 is highly expressed across cell types. Imputed RPL15 expression shows sub cell type structure introduced by the denoising process. Although the predictions show some sub cell type structure, they correlate better with the raw, than with the denoised observations. (Bottom) WFDC6 is not expressed in the dataset and imputation introduces low variation. The correlation of model predictions against raw observed all 0's can not be computed and correlation with the low level denoised expression signal is poor.

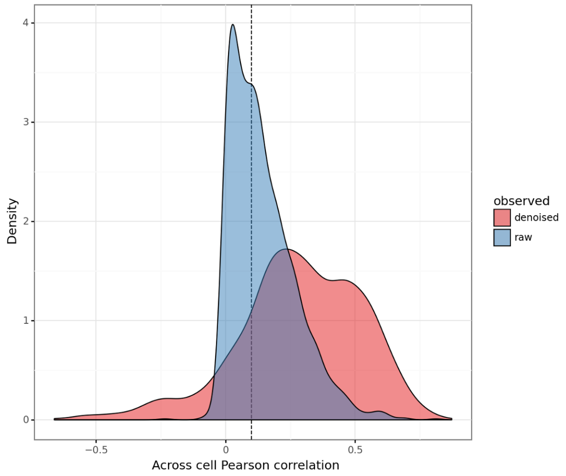


Supplementary Figure 4: Across cell Pearson correlation of test set genes when comparing model predictions against raw or denoised observed counts in test set genes. Dashed line indicates the empirical 0.1 Pearson correlation threshold for the across cell correlation of raw models beyond which genes are considered to capture a reasonable degree of cell specificity.

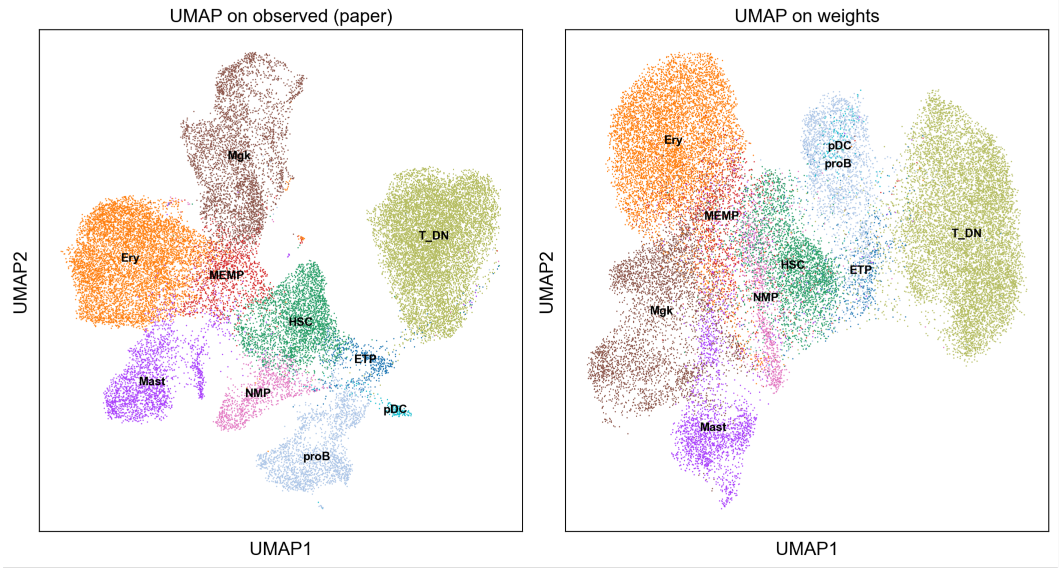


Supplementary Figure 5: UMAP embedding of the observed counts (left) and of the final layer weights of the single cell expression model for the HSC dataset. UMAP coordinates of the observed counts and cell annotations were provided in the public dataset (Park et al. 2020).

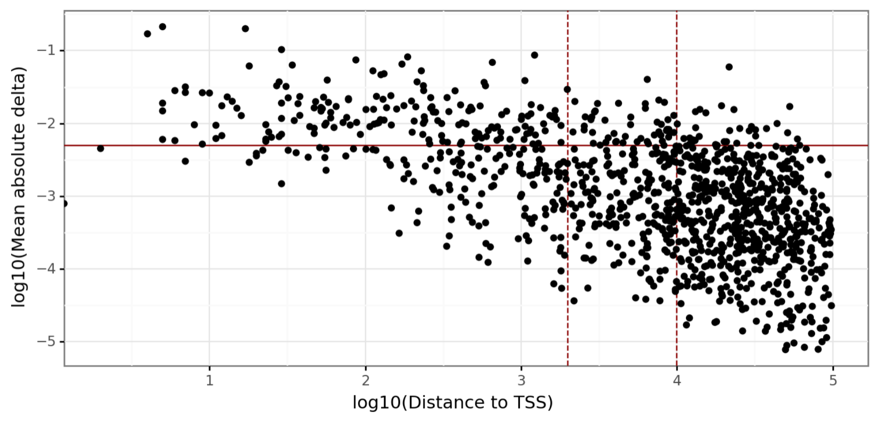


Supplementary Figure 6: Plotted is the log10 transformed absolute value of the delta prediction (variant – reference) for 1,154 whole blood eQTLs against the log10 transformed distance of the SNP to the canonical TSS of the linked gene. Dashed lines indicate 2 and 10 kb from the TSS, a rough guide for average promoter size and common sequences contexts of narrow models (e.g. scEP, NVMA). Solid line indicates an absolute mean delta of 0.005 an empirical threshold for single cell variant effect.

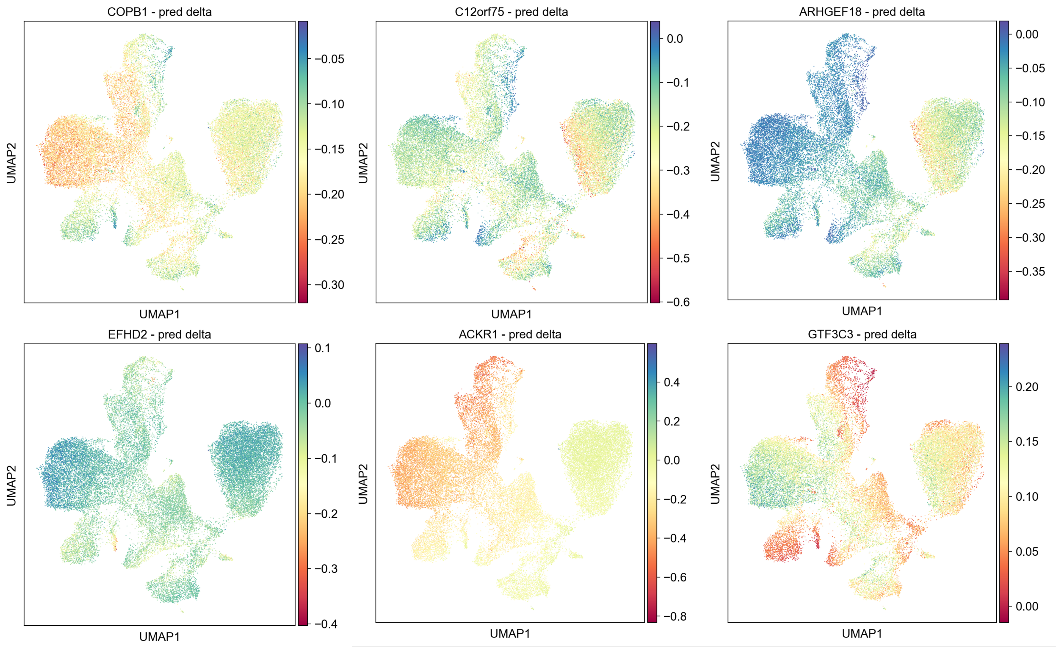


Supplementary Figure 7: Predicted effect of top six whole blood eQTL variants with high within cell type heterogeneity.

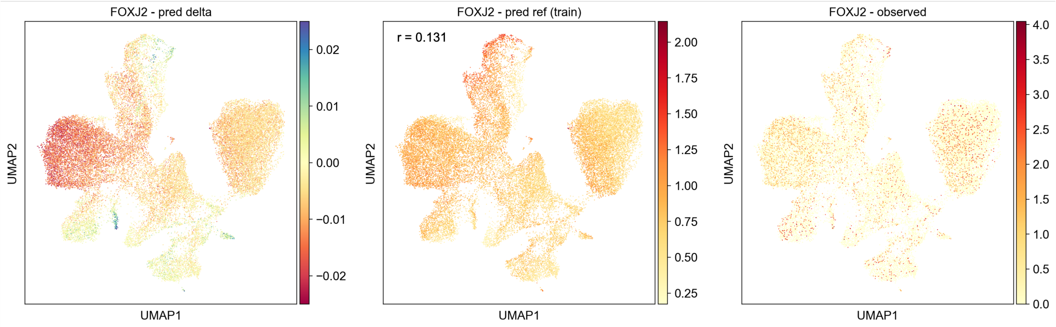


Supplementary Figure 8: Observed and predicted expression of FOXJ2 in the HSC dataset and the predicted effect (delta = variant – reference expression) of 12_8058041_A_G on FOXJ2 expression. The bivalent predicted effect is not explained by the expression of FOXJ2 itself.

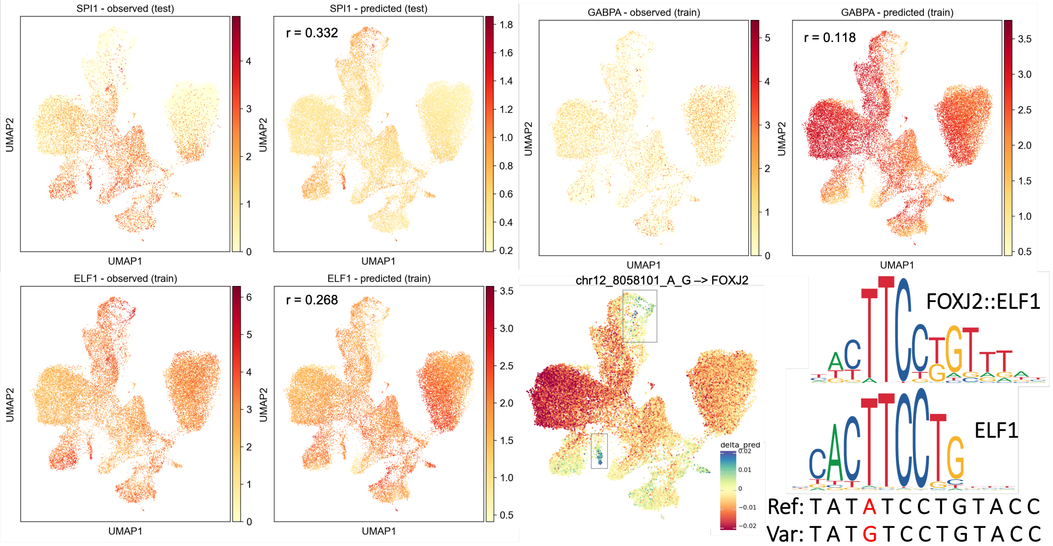


Supplementary Figure 9: The FOXJ2 eQTL (12_8058041_A_G) possibly disrupts an ETS-domain family binding site. The ETS-domain family of transcription factors consists of a multitude of transcription factors with a conserved binding motif. They can act as transcriptional activators or repressors, depending on the exact TF and co-regulating factors, one of which is FOXJ2. Exemplary expression in the HSC dataset shown for ETS-domain family members: SPI1 (PU.1), GABPA and ELF1. The positive predicted effect of 12_8058041_A_G on FOXJ2 in late Megakaryocytes (Mgk_late) and Eosinophils and Neutrophils (Eosin_Neutr) (black boxes) may be partly explained by a multi-faceted effect including enrichment of ELF1 (in Mgk_late & Eosin_Neutr) and SPI2 (in Eosin_Neutr) as well as depletion of GABPA (in Mgk_late and Eosin_Neutr). ELF1 motif was derived from JASPAR (MA0473.3 & MA1952)..

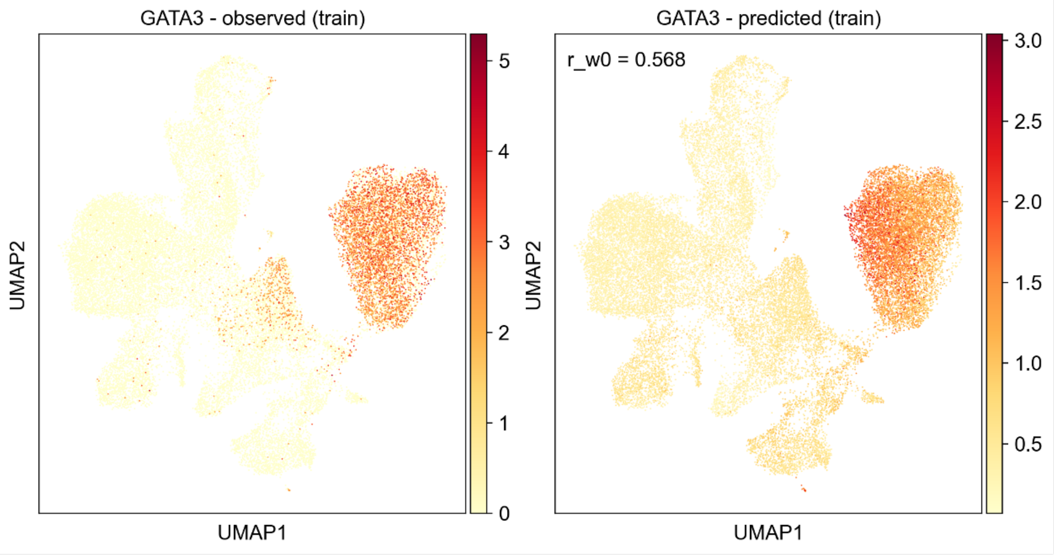


Supplementary Figure 10: The FOXJ2 eQTL (12_8058041_A_G) possibly disrupts a GATA binding site. GATA1 and GATA3 are expressed in Erythroid, HSC and T_DN cells where the variant is predicted to have a negative effect on transcription. As opposed to the positive effect it has on the whole blood level. Motifs derived from JASPAR (MA0035.4 & MA0037.3).

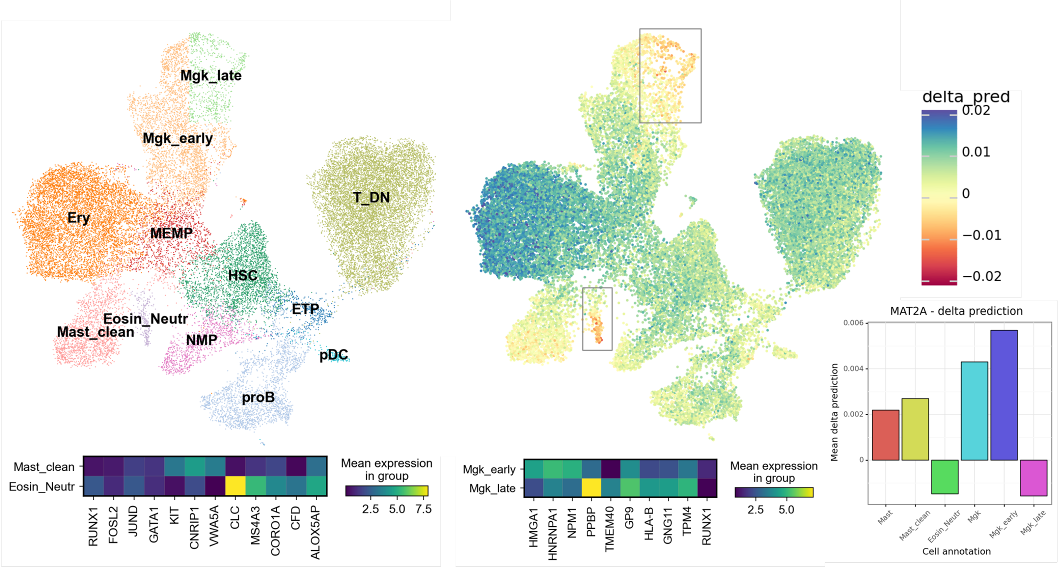


Supplementary Figure 11: Within cell type heterogeneity of an MAT2A linked whole blood eQTL (2_85501572_A_T). In eQTL studies the variant shows an expression reducing effect on MAT2A in whole blood and kidney and an enhancing effect in artery, prostate, colon and heart (GTEx portal). The variant also effects other genes: Increasing the expression of VAMP8 in brain and kidney and reducing it in the oesophagus and increasing the expression of GGCX in multiple tissue.

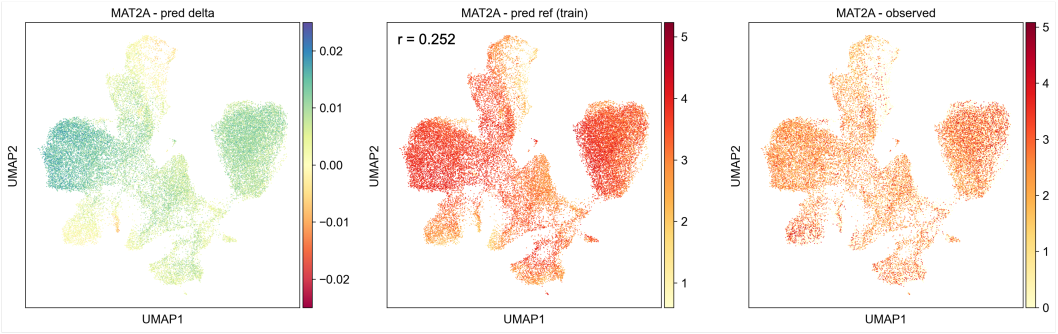


Supplementary Figure 12: Observed and predicted expression of MAT2A in the HSC dataset and the predicted effect (delta = variant – reference expression) of 2_85501572_A_T on MAT2A expression. The ambivalent predicted effect is not explained by the expression of MAT2A itself.

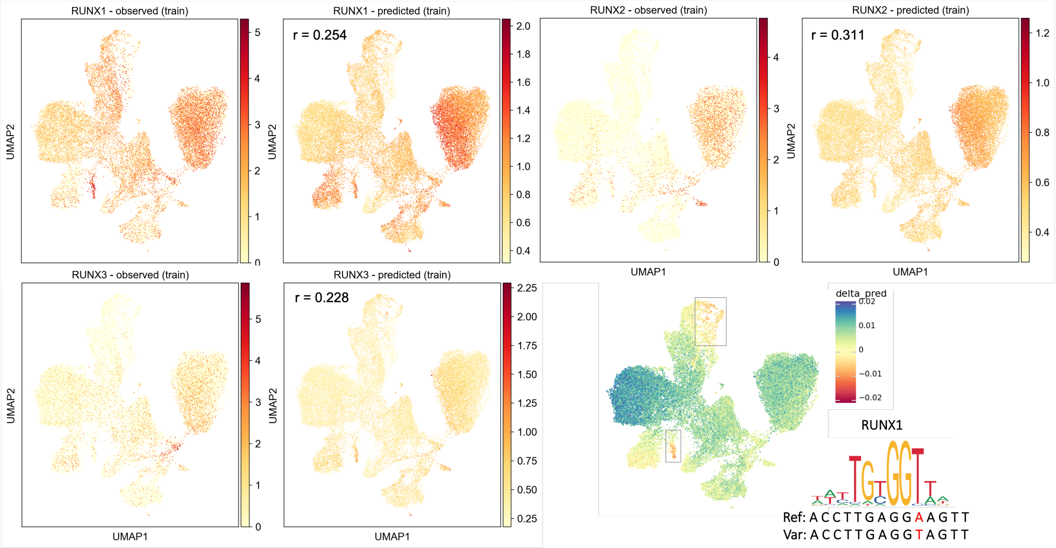


Supplementary Figure 13: A MAT2A eQTL (2_85501572_A_T) potentially enhances a binding motif for RUNX1 or other RUNX factors, the expression of which may explain the positive predicted effect on gene expression in the majority of cells in the HSC dataset. Motif derived from JASPAR (MA0002.1).

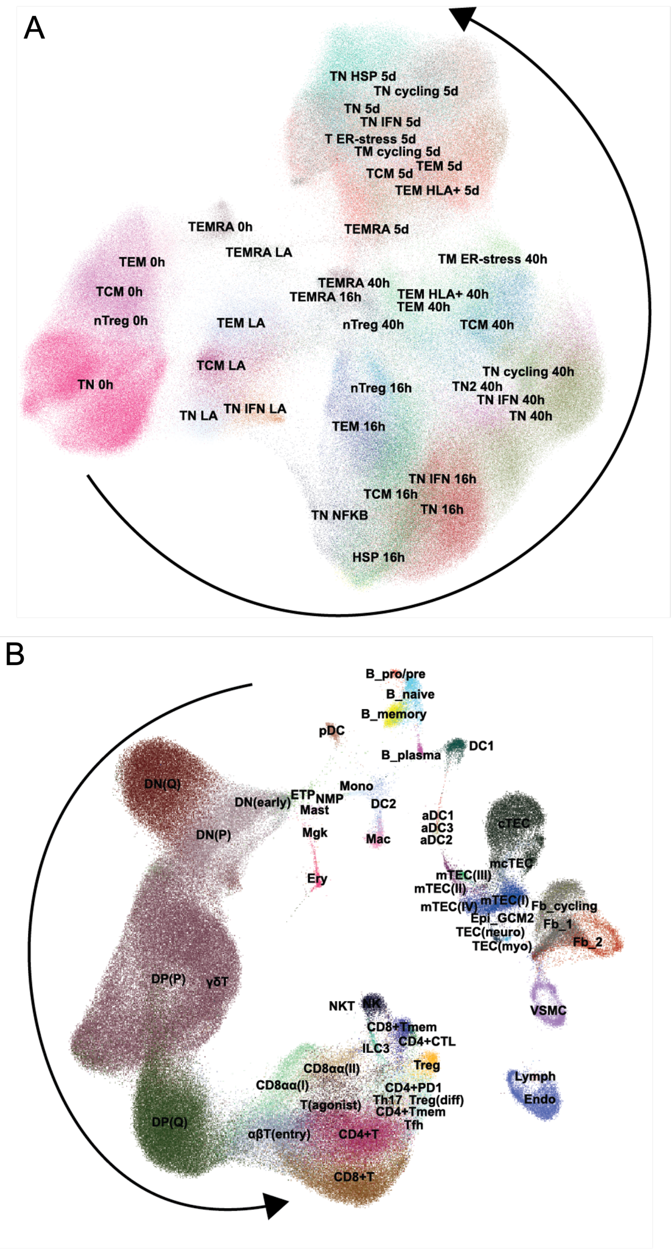


Supplementary Figure 14: Published UMAPs and cell annotations for (A) the CD4 T cell activation (Soskic et al. 2022) and (B) T cell development (Park et al. 2022) dataset. Black arrows are rough guides for the T cell activation trajectory from 0 h to 5 d after stimulation and for the T cell developmental trajectory from early thymic progenitors (ETP) to the fanning out of T cell types, respectively. Abbrevations adopted from the original publications (Park et al., Soskic et al.). A: ER - endoplasmic reticulum; IFN - cells expressing high levels of interferon; HLA - human leukocyte antigen; HSP - heat shock proteins; LA - lowly active / early activation stage; NF-$\kappa$B, nuclear factor $\kappa$B; nTreg - natural (i.e. thymus-derived) regulatory T cell; TCM - central memory T cell; TEM - effector memory T cell; TEMRA - effector memory T cells re-expressing CD45RA; TN - naive T cell. B: DC - dendritic cells; DN - double negative thymocyte; DP - double positive thymocyte; (Q) – quiescent; (P) – proliferating; ETP - early thymic progenitor; aDC - activated dendritic cells; pDC, plasmacytoid dendritic cells; Mono – monocyte; Mac – macrophage; Mgk – megakaryocyte; NK - natural killer cell; NKT -T lymphocytes co-expressing several NK cell-associated receptors; NMP - neutrophil-myeloid progenitor; Endo - endothelial cells; VSMC - vascular smooth muscle cells; Ery – erythrocytes; Epi – epithelial; Fb – fibroblast; ILC3 - Type 3 innate lymphoid cells; TEC – thymic epithelial cells; mTEC - medullar TEC; cTEC - cortical TECs; mcTEC - intermediate TEC population; Th17 - Th17-like cells.

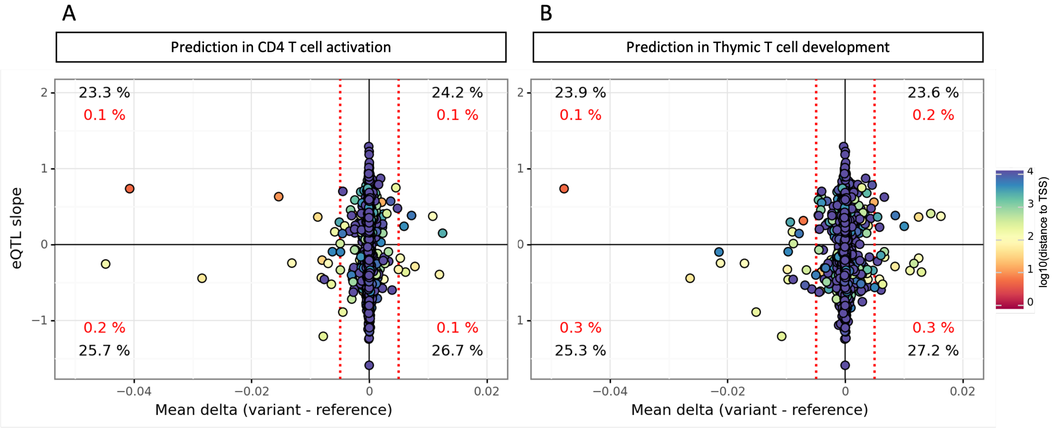


Supplementary Figure 15: Predicting the variant effect of 5,278 CD4 T cell activation eQTL associations and relating the average predicted variant effects across cells to the average slope of all significant associations per variant – eGene pair. Predictions were made using the using the CD4 T cell activation model (left) and T cell development model (right). Red dotted lines indicate our empirical variant effect threshold of +- 0.005. Indicated is the percentage of all associations (black) and all associations passing the empirical delta threshold (red) that fall within each respective quadrant indicating concordance or discordance between the average eQTL slope and the average predicted effect across cells.

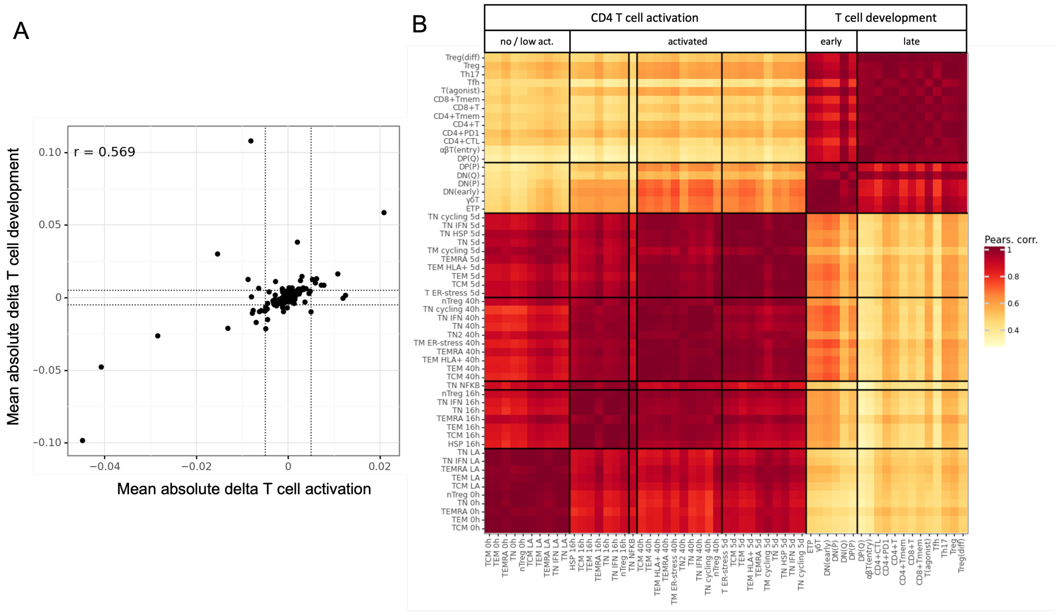


Supplementary Figure 16: A) Mean predicted variant effect across all cells of 5,278 eQTL variants on their linked eGenes, predicted using the CD4 T cell activation vs T cell development model. B) Pearson correlation of the mean predicted variant effects per cell type of 41 eQTL variants on their linked eGene where every variant has at least on per cell type effect larger than 0.005 or smaller than -0.005. Compared are annotated cell types from the CD4 T cell activation dataset (Soskic et al.) and the T cell development trajectory from the development dataset (Park et al.).

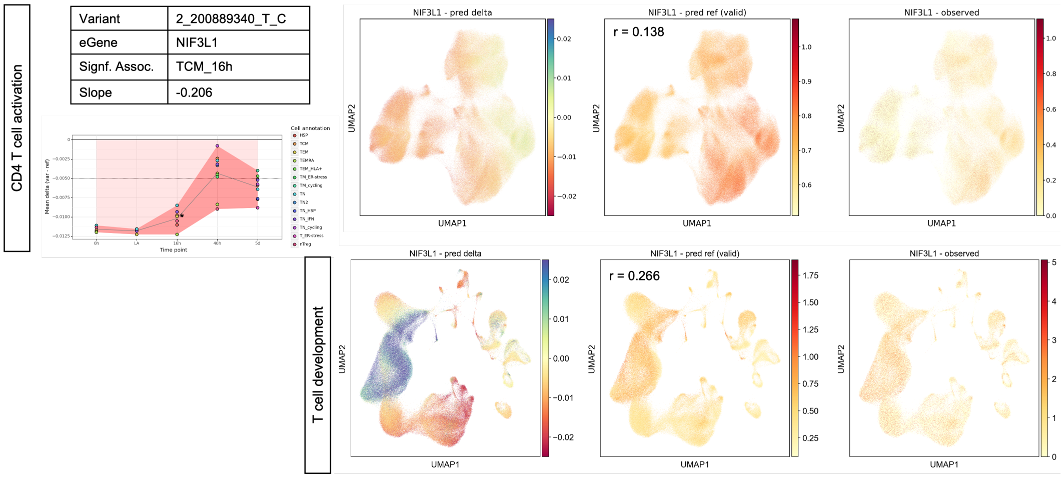


Supplementary Figure 17: Predicting the effect of the eQTL variant 2_200889340_T_C on its eGene NIF3L in CD4 T cell activation and T cell development. The eQTL was identified with a significant effect only in central memory T cells (TCM) 16h after stimulation with a negative slope (less expression in variant). We plotted the average predicted delta (var – ref) in NIF3L expression for each cell population along the time course of stimulation. The dashed line indicates the -0.005 emprical variant effect threshold. The asterisk marks the TCM_16h cell population. Plotting the predicted effect per cell in T cell development, reveals an effect with different directionality: A negative delta in quiescent double negative (DN) and double positive (DP) thymocytes, in more mature T cells as well as in B cells, dendritic cells and monocytes, but a positive delta in proliferating DN and DP thymocytes as well as erythroid cells, megakaryocytes and fibtroblasts (compare to Supplementary Figure 14). NIF3L1 was in the validation set for model training.

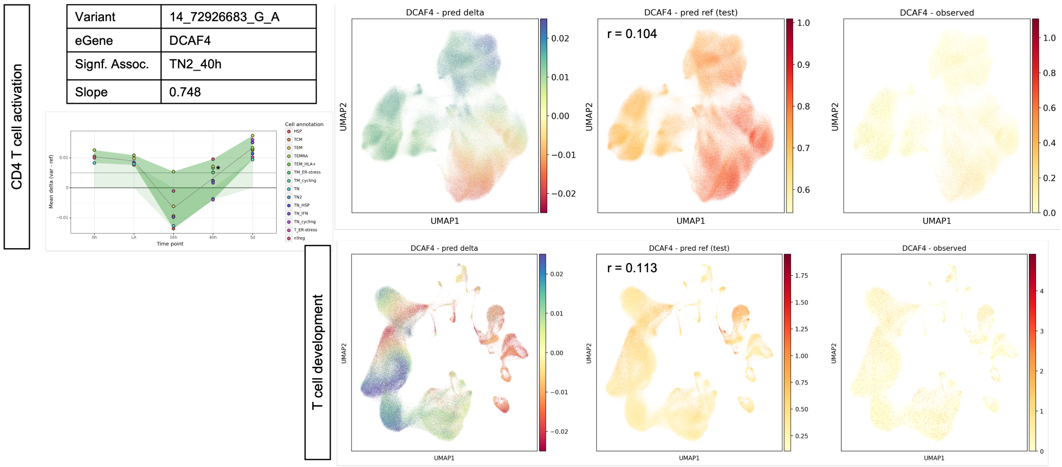


Supplementary Figure 18: Predicting the effect of the eQTL variant 14_72926683_G_A on its eGene DCAF4 in CD4 T cell activation and T cell development. The eQTL was identified with a significant effect only in a subset of naïve T cells (TN2) 16h after stimulation with a positive slope (more in expression in variant). We plotted the average predicted delta (var – ref) in DCAF4 expression for each cell population along the time course of stimulation. The dashed line indicates the 0.005 empirical variant effect threshold. The asterisk marks the TCM_16h cell population. Plotting the predicted effect per cell in the T cell development dataset indicates a negative effect in the majority of non T cell cell types. Moreover, the predicted effect is negative in early T cell progenitors but is positive in quiescent double negative thymocytes and in double positive (DP) thymocytes further along the trajectory including late proliferating DPs and quiescent DPs as well as more mature T cells. Interestingly, the predicted effect direction shifts within the annotated DP proliferating cell type, suggesting a finer grained annotation may be revealed by the variant effect (compare to Supplementary Figure 14). DCAF4 was in the test set for model training.

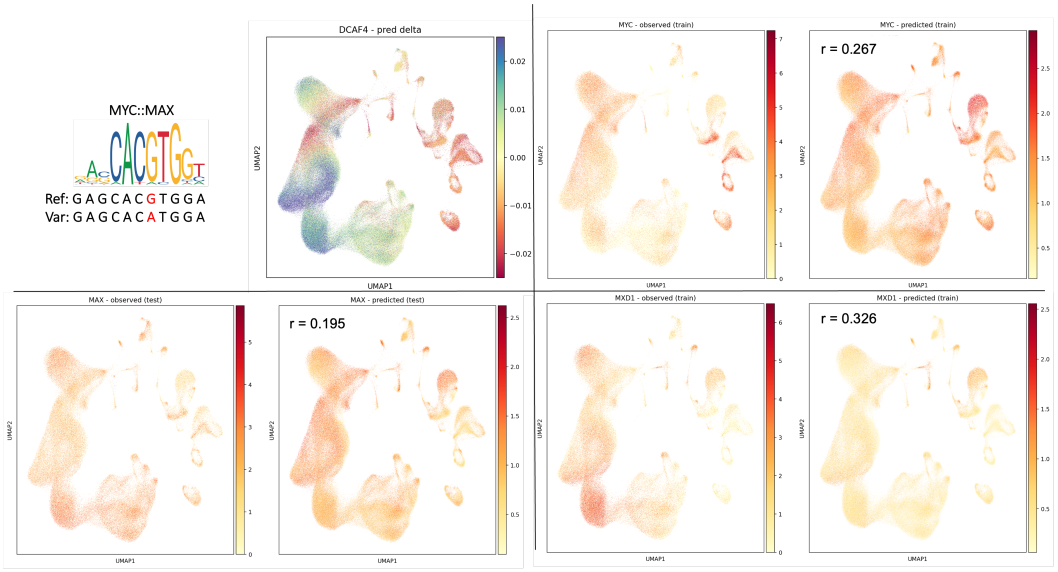


Supplementary Figure 19: Comparing the predicted effect of the DCAF4 eQTL 14_72926683_G_A with the observed and predicted expression of MAX, MYC and MXD1 (Mad1) in T cell development. The variant disrupts an E-box motif . The relative change of observed and predicted expression of MYC and MXD1 along the T cell development trajectory may explain the different direction of the predicted variant effect. Myc-Max heterodimers act as transcriptional activators, whereas Mad::Max, with identical DNA sequence preferences, act as transcriptional suppressors. Binding motif were derived from JASPAR (MA0059.1 & MA0058.2).

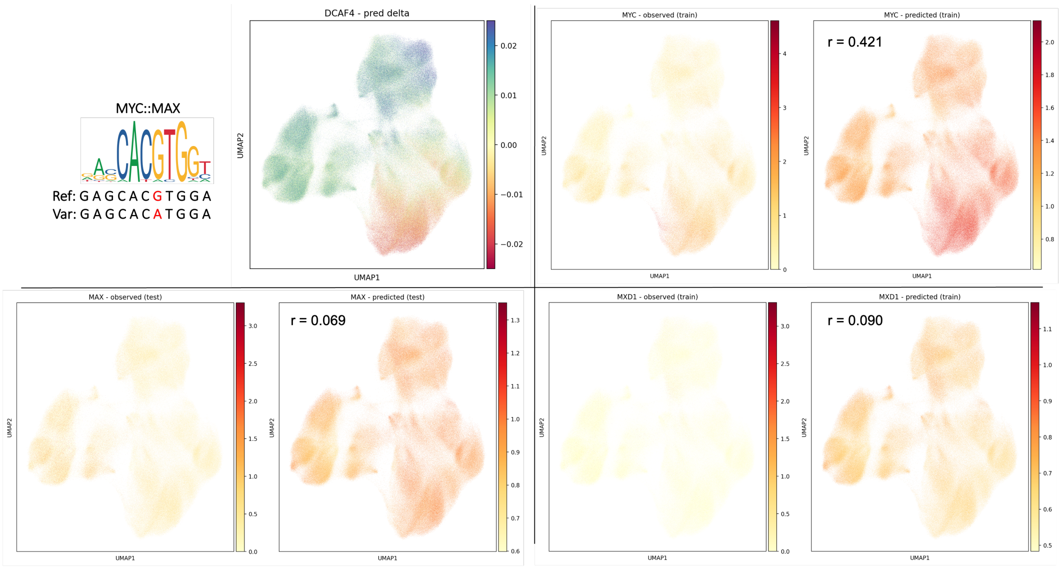


Supplementary Figure 20: Comparing the predicted effect of the DCAF4 eQTL 14_72926683_G_A with the observed and predicted expression of MAX, MYC and MXD1 (Mad1) in T cell activation. The variant disrupts an E-box motif. The predicted negative (reducing expression) effect roughly tracks the high expression of MYC in activated T cells 16 and 40 h after stimulation. Myc-max heterodimers act as transcriptional suppressors. The positive predicted impact (enhancing expression) fits to cells with lower levels of MYC expression and higher levels of observed MAX expression although MAX expression itself was not well captured by the model. MXD1 expression is very sparse in the T cell activation dataset.Binding motif of MYC::MAX was derived from JASPAR (MA0059.1).

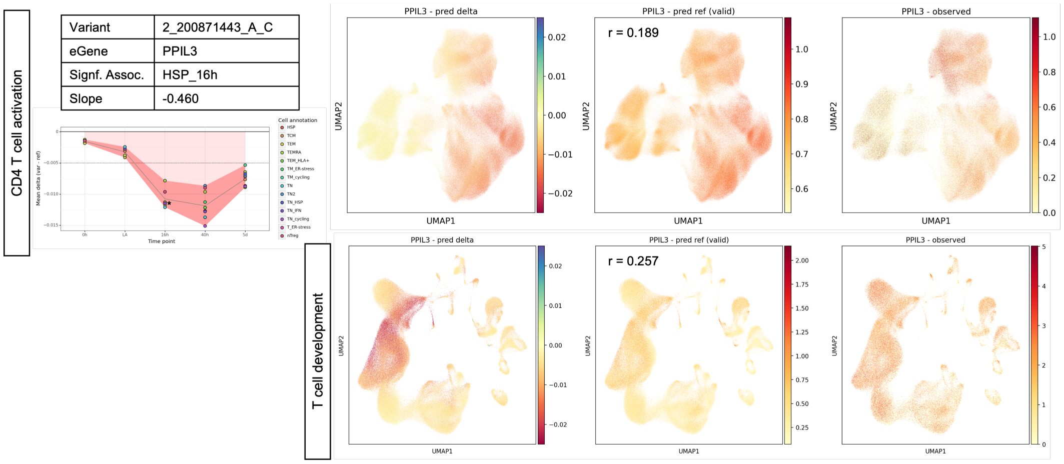


Supplementary Figure 21: Predicting the effect of the eQTL variant 2_200871443_A_C on its eGene PPIL3 in CD4 T cell activation and T cell development. The eQTL was identified with a significant effect only in cells with high levels of heat shock proteins (HSP) 16 h after stimulation, with a negative slope (less expression in variant). We plotted the average predicted delta (var – ref) in PPIL3 expression for each cell population along the time course of stimulation. The dashed line indicates the -0.005 empirical variant effect threshold. The asterisk marks the HSP_16h cell population. In T cell activation, the predicted effect is strongest in activated T cells , 16 and 40 h after stimulation. In T cell development the effect is strongest in proliferating double negative and double positive thymocytes (compare to Supplementary Figure 14). PPIL3 was in the validation set for model training.

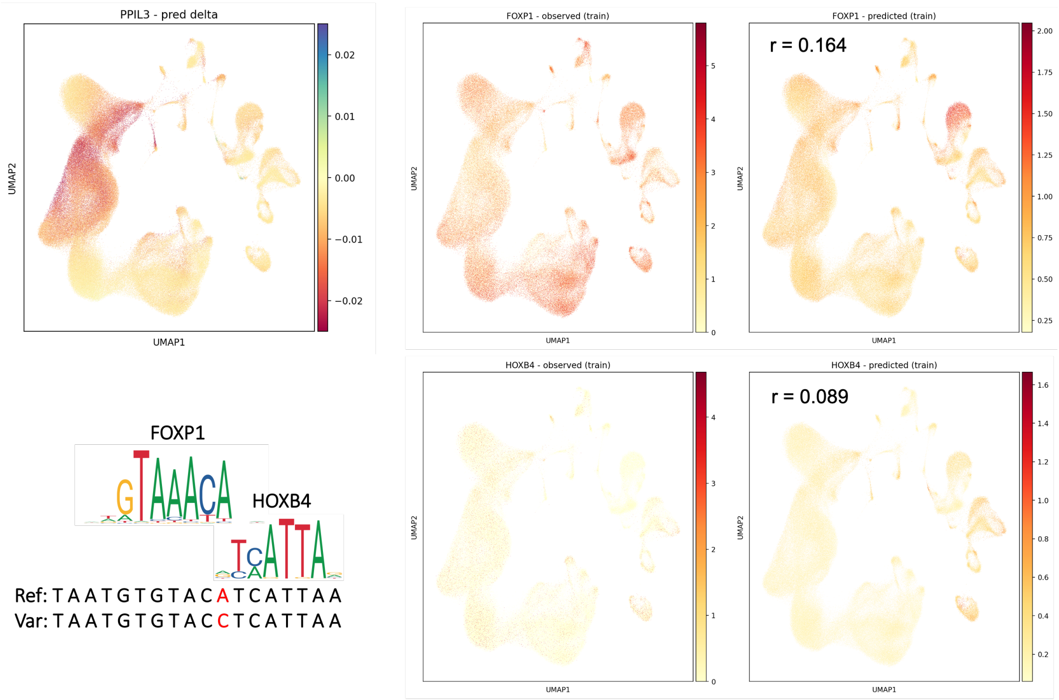


Supplementary Figure 22: Comparing the predicted effect of the PPIL3 eQTL 2_200871443_A_C with the observed and predicted expression of FOXP1 and HOXB4. Note that a plethora of transcription factors may bind to FOX-like and HOX-like motifs and FOXP1 and HOXB4 were used as examples. Motifs for FOXP1 (MA0481.1) and HOXB4 (MA1499.1) were derived from JASPAR.
